# A genetically encoded fluorescent sensor enables sensitive and specific detection of IDH mutant associated oncometabolite D-2-hydroxyglutarate

**DOI:** 10.1101/2024.09.25.615072

**Authors:** Kristian A. Choate, Wren W.L. Konickson, Matthew J. Jennings, Paul B. Mann, Robert J. Winn, David O. Kamson, Evan P.S. Pratt

**Affiliations:** Department of Biology, Northern Michigan University, Marquette, MI, 49855, USA; Upper Michigan Brain Tumor Center, Northern Michigan University, Marquette, MI, 49855, USA; Department of Chemistry, Northern Michigan University, Marquette, MI, 49855, USA; School of Clinical Sciences, Northern Michigan University, Marquette, MI, 49855, USA; Department of Oncology, Johns Hopkins University School of Medicine, Baltimore, MD, 21287, USA; Department of Neurology, Johns Hopkins University School of Medicine, Baltimore, MD, 21287, USA

**Keywords:** Fluorescence Resonance Energy Transfer, Fluorescent Protein, Genetically Encoded, D-2-Hydroxyglutarate, Isocitrate Dehydrogenase, Glioma

## Abstract

D-2-hydroxyglutarate (D-2-HG) is an oncometabolite that accumulates due to mutations in isocitrate dehydrogenase 1 and 2 (*IDH1/2*). D-2-HG may be used as a surrogate marker for *IDH1/2* mutant cancers, yet simple and specific methods for D-2-HG detection are limited. Here, we present the development and characterization of a genetically encoded fluorescent sensor of D-2-HG (D2HGlo). D2HGlo responds to clinically relevant concentrations of D-2-HG, demonstrates exceptional selectivity and can quantify D-2-HG in various body fluids and glioma tumor supernatants. Additionally, analysis of tumor lysates using D2HGlo accurately predicted the IDH mutational status of gliomas. Collectively, these results suggest that D2HGlo may be an *in vitro* diagnostic device for the detection and monitoring of IDH mutant cancers through liquid biopsies. In addition to D2HGlo’s clinical utility, we also present preliminary findings for its adaptation to the cellular environment. To assess D-2-HG production in living immortalized glioma cells, we engineered D2HGlo sensors that localize to subcellular compartments, which yielded findings of elevated D-2-HG in the cytosol, mitochondria, and nucleus of *IDH1* mutant cells.

## Introduction

D-2-hydroxyglutarate (D-2-HG) is an oncometabolite that is associated with several forms of cancer, including glioma and acute myeloid leukemia (AML)^1,2^. Cancers that produce elevated levels of D-2-HG harbor mutations that replace the canonical arginine with another amino acid in the active site of the enzyme isocitrate dehydrogenase (IDH). The function of IDH is to convert isocitrate to α-ketoglutarate, reducing NADP^+^ to NADPH. Neomorphic mutations at positions R132 in *IDH1* and R140 or R172 in *IDH2* cause both enzymes to catalyze the reduction of α-ketoglutarate to D-2-HG and have been shown to be an early event in the development of cancer^3^. The accumulation of D-2-HG promotes oncogenesis through the competitive inhibition of enzymes belonging to the α-ketoglutarate/Fe(II)-dependent dioxygenase family^4^. This leads to epigenetic alterations associated with the Global-CpG Island Methylation Phenotype (G-CIMP) and aberrant expression of oncogenes and tumor suppressor genes^5^. Cells associated with G-CIMP are thought to be less aggressive in part due to the delayed repair of double stranded DNA breaks. Resultantly, *IDH1/2* mutant cells are highly sensitive to DNA damaging agents such as temozolomide or radiation and are more susceptible to poly(ADP)-ribose polymerase (PARP) inhibitors. In general, IDH mutations are associated with a better patient outcome, as they render cells more vulnerable to death^6^ and demonstrate reduced levels of migration, angiogenesis, and invasion^7^; thus, a mutant version of this enzyme is a significant and positive prognostic biomarker^8^.

Given the considerable impact of *IDH* mutations on cancer phenotype and patient prognosis, efficient and timely identification of this mutation is critical. Due to the lengthy analytical times of currently available diagnostic methods such as sequencing and immunohistochemistry (IHC), clinicians do not have knowledge of the tumor’s genetic characteristics until days or weeks post-surgery. As an alternative to the direct detection of *IDH* mutations, D-2-HG may be utilized as a rapid fluid-based surrogate marker^1,9–14^. Current methods for detecting D-2-HG include gas chromatography-mass spectrometry (GC-MS), liquid chromatography-tandem mass spectrometry (LC-MS/MS) and magnetic resonance spectroscopy (MRS)^15–18^. While these methods are well characterized, they are not amenable for rapid and routine use in a clinical laboratory. Studies which have explored the relationship between D-2-HG level and disease largely disagree on optimal sample type and the relative quantity of oncometabolite corresponding to *IDH* mutational status (Table 1). This lack of consensus warrants a standardized method for the noninvasive, rapid quantification of D-2-HG in body fluids to allow for the preoperative, intraoperative, and/or postoperative detection of an *IDH* mutation. Additionally, the recent success of Vorasidenib, a pan mutant IDH inhibitor (INDIGO trial), makes real-time monitoring of D-2-HG levels in patients imperative as its depletion can be utilized to gauge the effectiveness of inhibitor therapy^19^.

**Table 1.**
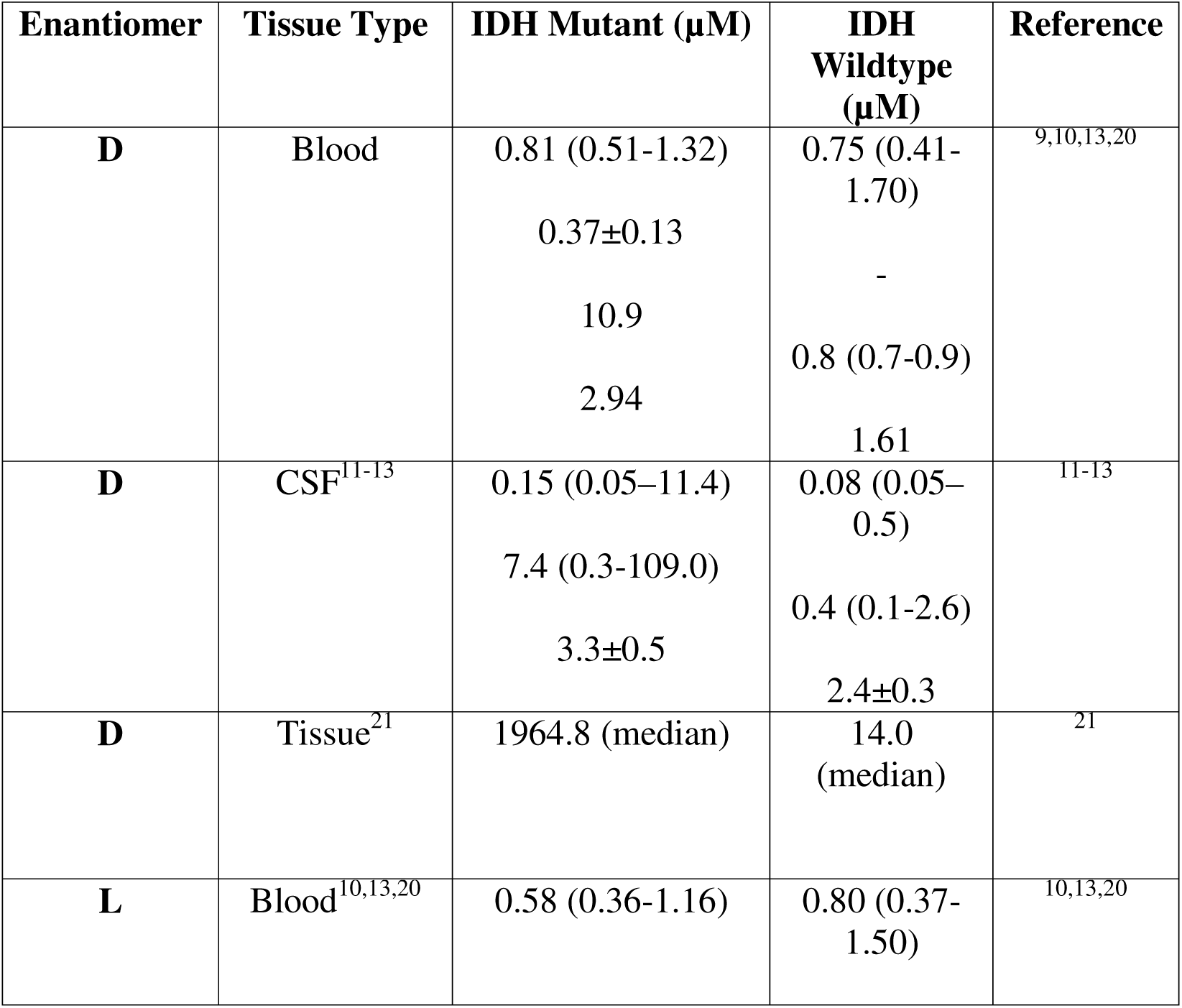
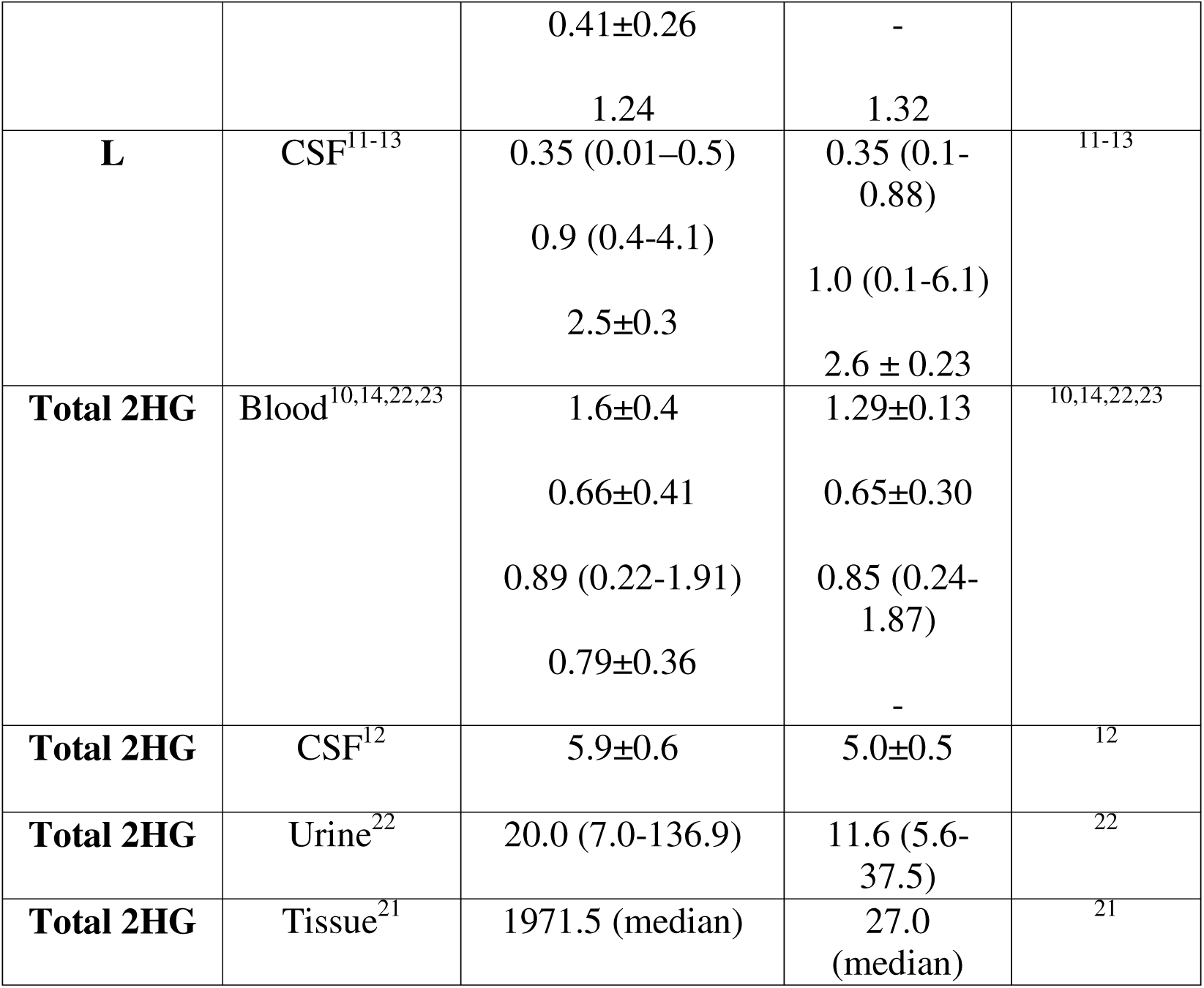
Summary of D-2-HG or L-2-HG concentrations in biological matrices as presented in current literature.

In this study, we present a genetically encoded fluorescent sensor that is highly specific for D-2-HG compared to other structurally similar metabolites and demonstrates a greater affinity for D-2-HG over L-2-HG. D2HGlo robustly quantifies D-2-HG in a variety of biological fluids and accurately predicts the IDH mutational status when using glioma tumor supernatants. The reportable range of detection for D2HGlo suggests that it may be a powerful tool for detecting elevated levels of D-2-HG, measuring the efficacy of pharmaceutical inhibitors, and monitoring remission vs. recurrence in patients with IDH mutant cancers. This sensor also facilitated preliminary investigations of the intracellular distribution of D-2-HG in living human cells to include its presence in the nuclear compartment.

## Results

### Development of a genetically encoded fluorescent sensor for D-2-hydroxyglutarate

The transcription factor D-2-HG dehydrogenase regulator (DhdR) was recently identified as a D-2-HG-binding protein and key mediator of D-2-HG catabolism in *Achromobacter denitrificans*, NBRC 15125^24^. To engineer a genetically encoded fluorescent sensor of D-2-HG, we leveraged DhdR as a D-2-HG binding domain. We designed a fluorescence resonance energy transfer (FRET) based sensor by inserting DhdR between the fluorescent proteins Enhanced Cyan Fluorescent Protein (ECFP) and a circularly permuted variant of Venus (cpV173). We hypothesized that D-2-HG binding DhdR would elicit a conformational change, resulting in increased FRET between the two fluorescent proteins (Fig. 1a). To test this prediction, we purified the DhdR-based fluorescent sensor and exposed the sensor to increasing concentrations of D-2-HG, ranging from 10 nM to 1 mM. The FRET ratio was determined by exciting the purified protein at 440 nm and dividing the fluorescent signal collected at 531 nm by the fluorescent signal at 485 nm. We found that at concentrations greater than 100 nM D-2-HG, the FRET ratio increased in a dose-dependent manner until plateauing at approximately 100 µM (Fig. 1b). The apparent binding affinity (K_d_’) of this sensor for D-2-HG was 4.00 ± 1.64 µM with an estimated linear range of 1 µM-50 µM. The dynamic range, defined as the maximum FRET ratio (R_max_) divided by the minimum FRET ratio (R_min_), was 1.37 ± 0.01.

**Figure 1.**
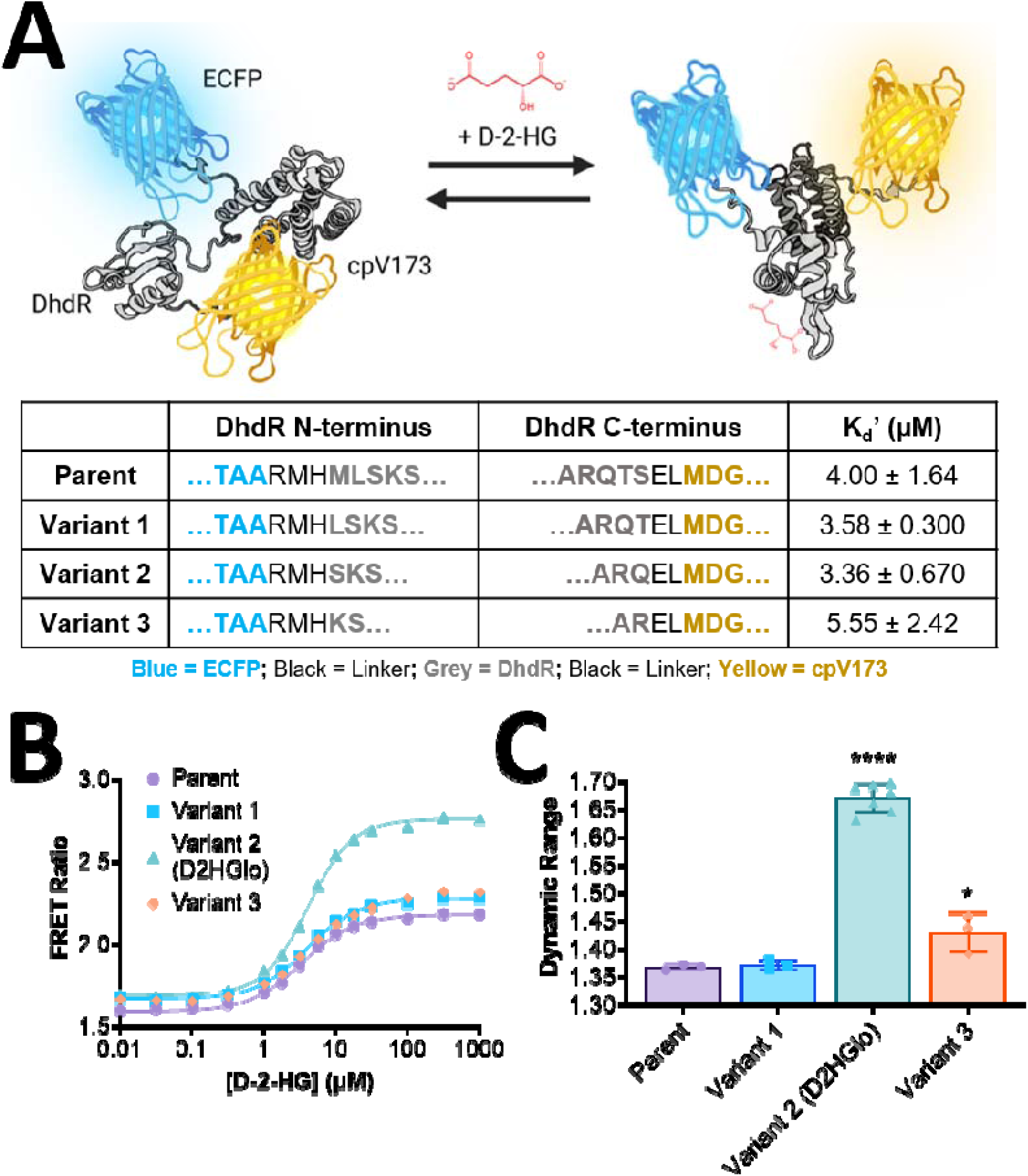
Design and optimization of a genetically encoded fluorescent sensor of D-2-hydroxyglutarate (D-2-HG). (A) Schematic representation of a FRET-based D-2-HG sensor. The transcription factor DhdR is flanked by two fluorescent proteins (ECFP and cpVenus173) that act as a FRET pair. The structure of DhdR was predicted using AlphaFold and the FRET sensor model was generated in BioRender. The amino acid sequences at the N-terminus and C-terminus of DhdR are shown for the parent sensor and three sensor variants, along with the average K_d_’ ± standard deviation for each construct. (B) Representative D-2-HG binding curves for the parent FRET sensor and three sensor variants. Log [D-2-HG] is plotted against the FRET ratio for each construct. (C) Comparison of the dynamic range (R_max_/R_min_) of the parent sensor and three sensor variants. The average dynamic range ± standard deviation is shown for 3-7 independent experiments. Statistical analysis was performed using a one-way ANOVA test with *post hoc* Tukey (****, P < 0.0001 compared with Parent, Variant 1 and Variant 3; *, P < 0.05 compared with Parent and Variant 1).

The binding properties of the initial FRET sensor suggested that it was suited to detect D-2-HG in the reported physiological range (Table 1). However, the dynamic range was less than other FRET-based sensors developed for similar metabolites^25,26^. Thus, to improve the dynamic range, we manipulated the linker regions connecting DhdR with ECFP and cpV173. We generated three sensor variants by systematically removing one amino acid at a time from both the N-terminus and C-terminus of DhdR. While the K_d_’ for D-2-HG remained relatively unchanged among the four sensors (Fig. 1b), Variant 2 exhibited a significantly greater dynamic range compared with the parent sensor and other two variants (Fig. 1c). We named this high-performing variant D2HGlo. D2HGlo has a K_d_’ of 3.36 ± 0.67 µM for D-2-HG and an *in vitro* dynamic range of 1.67 ± 0.03. Emission spectra of D2HGlo that were collected in the presence of increasing concentrations of D-2-HG revealed that the rise in FRET ratio was due to a drop in fluorescent signal at 485 nm and concomitant increase in the fluorescent signal at 531 nm (Supplemental Fig. 1). Using FRET ratios obtained for lower concentrations of D-2-HG ranging from 50 – 500 nM (Supplemental Fig. 2), we calculated the limit of detection to be 308 nM using the equation LOD= 3σ/S, where σ is the standard deviation of the blank and S is the slope of the standard curve.

### *In vitro* characterization of purified D2HGlo

We next sought to characterize the specificity of D2HGlo by evaluating its response to six Citric Acid Cycle intermediates and three additional metabolites that structurally resemble D-2-HG (pyruvate, lactate, and glutamate). D2HGlo did not respond to any of the metabolites at concentrations up to 100 µM (Fig. 2b and Supplemental Fig. 3). These results indicate that D2HGlo has exceptional specificity for D-2-HG compared to other relevant biological molecules. We then evaluated the effect of pH and temperature on D2HGlo *in vitro*. pH did not drastically alter the sensor’s affinity for D-2-HG (Fig. 2c, Supplemental Table 1); however, the dynamic range of D2HGlo was decreased at pH 6.5 and increased at pH 8 relative to measurements performed at pH 7.4. Lastly, we assessed if incubating the sensor with ligand at increased temperatures of 30°C or 37°C impacted the ability of D2HGlo to reliably quantify D-2-HG in comparison to experiments performed at room temperature. We found no significant difference in K_d_’ or dynamic range when comparing room temperature and 30°C or 37°C incubations (Fig. 2d, Supplemental Table 1). Thus, D2HGlo specifically detects D-2-HG over normally occurring metabolites at physiological pH and at temperatures amenable for *in vitro* or *in situ* studies.

**Figure 2.**
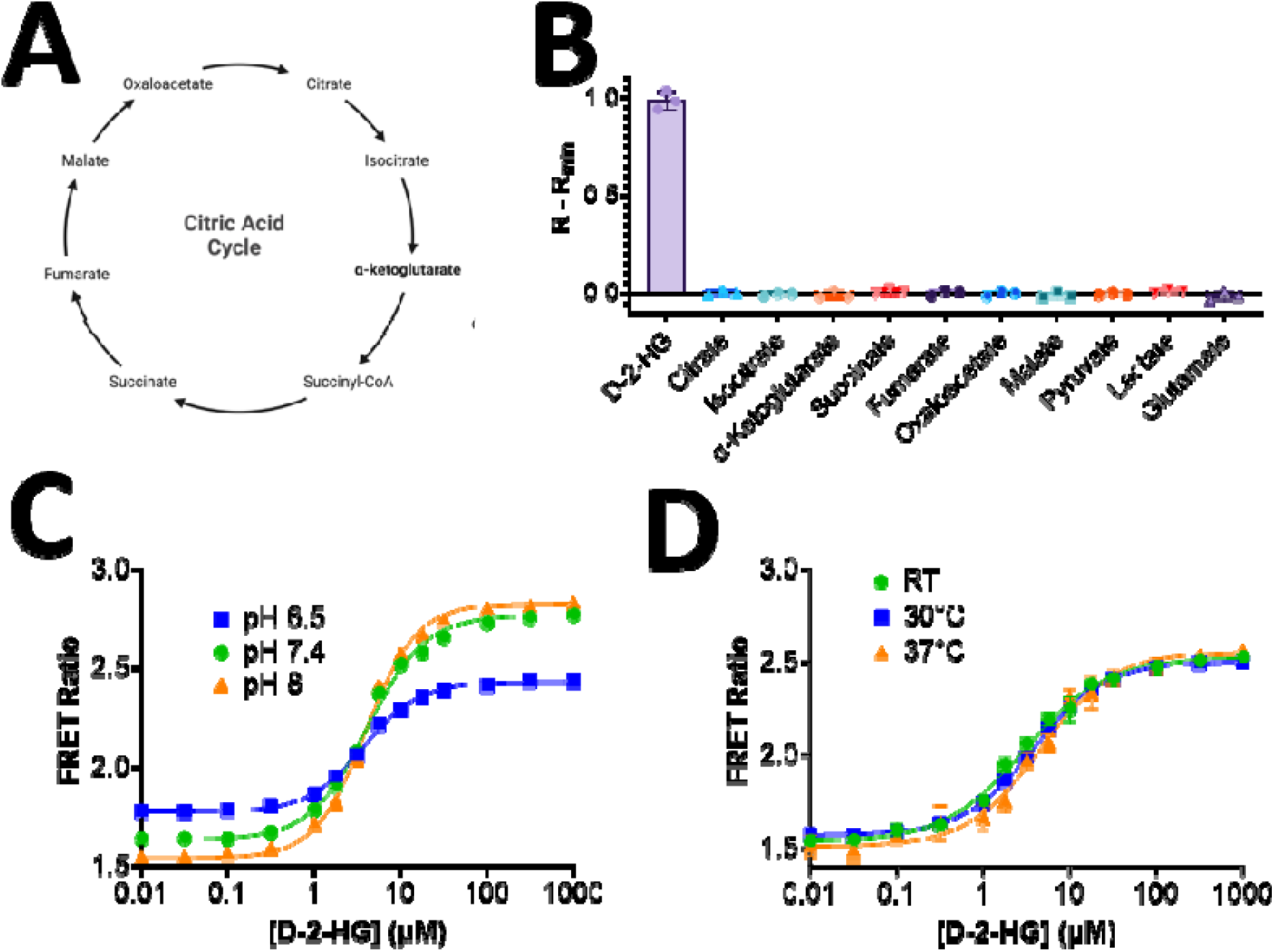
*In vitro* characterization of purified D2HGlo. (A) Diagram of the Citric Acid Cycle, highlighting key intermediates. (B) Selectivity of D2HGlo for D-2-HG compared with seven Citric Acid Cycle intermediates (citrate, isocitrate, α-ketoglutarate, succinate, fumarate, oxaloacetate and malate). In addition, pyruvate, lactate and glutamate were tested. R-R_min_ was calculated by subtracting the FRET ratio in the presence of 100 nM metabolite (R_min_) from the FRET ratio in the presence of 100 µM metabolite (R). The average ± standard deviation for each condition is shown for three independent experiments. (C) The influence of pH on *in vitro* fluorescence measurements. Purified D2HGlo was diluted in three different buffers (pH 6.5, pH 7.4 and pH 8), and the representative binding curves are shown.(D) The impact of temperature on *in vitro* fluorescence measurements. D2HGlo was incubated at 30°C or 37°C in the presence of D-2-HG for 15 minutes prior to fluorescent measurements and the representative binding curves are shown.

### *In vitro* characterization of the response of D2HGlo to L-2-HG

The excess production of D-2-HG in human cancers occurs due to the neomorphic enzymatic activity of mutant IDH1/2, whereas L-2-HG is a product of the promiscuous activity of lactate dehydrogenase and malate dehydrogenase (Fig. 3a)^27,28^. For D2HGlo to be considered a robust analytical tool for measuring D-2-HG in biological samples, discrimination between D-2-HG and L-2-HG is essential. Therefore, we directly compared D-2-HG and L-2-HG binding to D2HGlo *in vitro* (Fig. 3b). We found that the K_d_’ of D2HGlo for L-2-HG was 34.3 ± 4.95 µM, compared with a K_d_’ of 3.52 ± 0.84 µM for D-2-HG. Thus, D2HGlo has a greater affinity for D-2-HG compared with its enantiomer, L-2-HG. Since the physiologically normal levels of L-2-HG in human body fluids are approximately 0.1-6.1 µM (Table 1), we selected a concentration within that range (1 µM) and exceeding that range (10 µM). We found that at 1 µM, L-2-HG elicited a 1.43% increase in the FRET ratio above baseline levels, compared with an 8.22% increase in response to the same concentration of D-2-HG. 10 µM L-2-HG elicited a 9.58% increase, in contrast to the 47.3% increase elicited by the same concentration of D-2-HG (Fig. 3c). While 10 µM L-2-HG did induce a response, concentrations of L-2-HG above this level have rarely been reported in body fluids of individuals without an oncogenic IDH mutation or metabolic disorder such as L-2-hydroxyglutaric aciduria.

**Figure 3.**
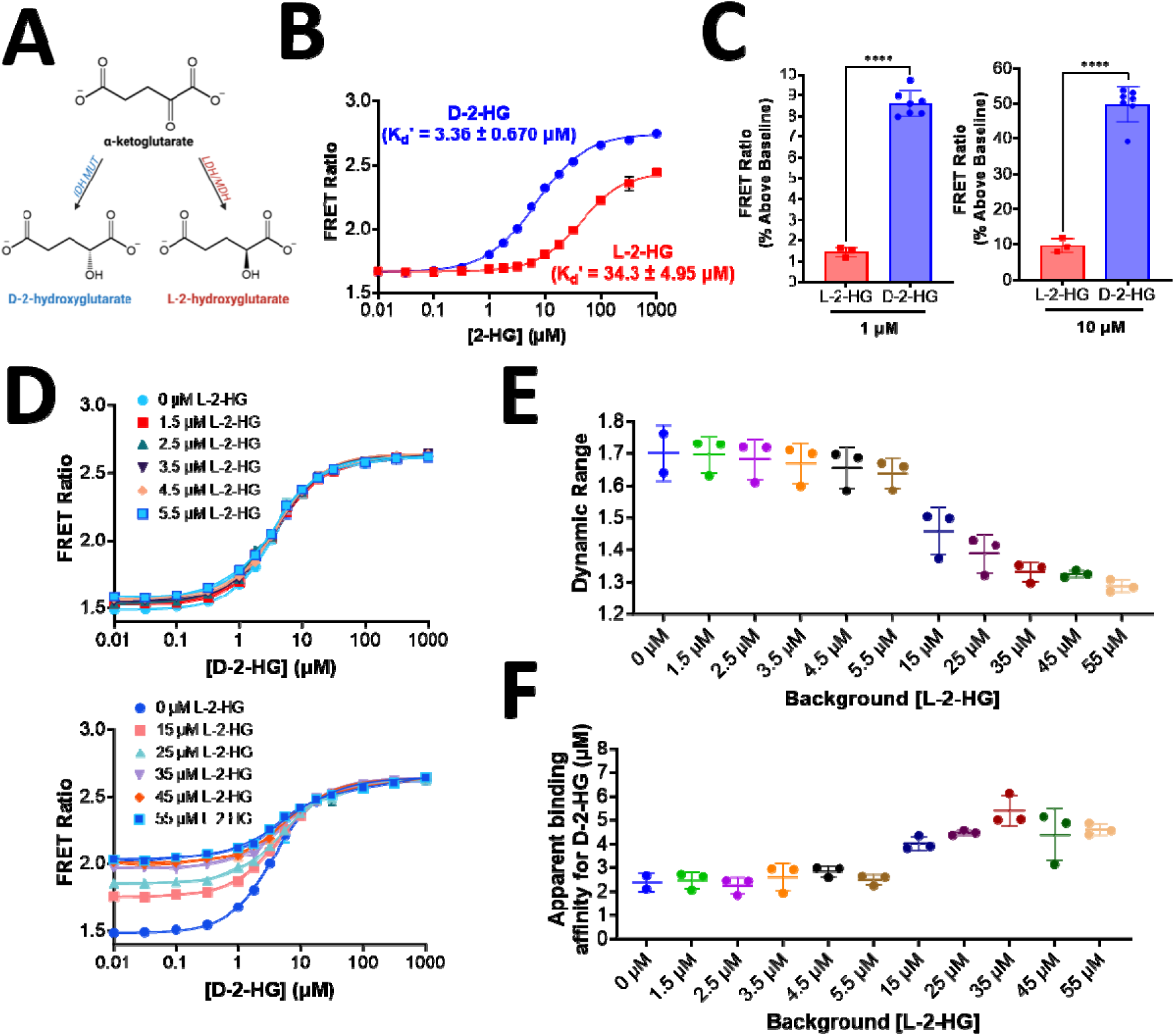
*In vitro* characterization of the response of D2HGlo to L-2-HG. (A) IDH1-R132H catalyzes the conversion of α-ketoglutarate to D-2-HG. L-2-HG is produced by lactate dehydrogenase (LDH) and malate dehydrogenase (MDH). (B) Representative *in vitro* D-2-HG and L-2-HG binding curves for purified D2HGlo. The average K_d_’ ± standard deviation is shown for D-2-HG (blue) and L-2-HG (red) and represents 3-7 independent experiments. (C) The percent increase in FRET ratio above baseline levels in response to D-2-HG and L-2-HG at 1 µM (left) and 10 µM (right). The average ± standard deviation for each condition is shown for three independent experiments. Statistical analysis was performed using an unpaired t-test (****, P < 0.0001). (D) Representative *in vitro* D-2-HG binding curves for purified D2HGlo in a background of low concentrations of L-2-HG (1.5 – 5.5 µM, top) or high background concentrations of L-2-HG (15 – 55 µM, bottom). (E) The *in vitro* dynamic range of D2HGlo determined from three independent D-2-HG titrations in the presence of background L-2-HG ranging from 1.5 – 55 µM. (F) The K_d_’ of D2HGlo for D-2-HG from three independent experiments performed in the presence of L-2-HG at concentrations ranging from 1.5 – 55 µM.

To further assess the impact of a physiologically relevant concentration of L-2-HG on the ability of D2HGlo to quantify D-2-HG, we performed D-2-HG titrations in a background of L-2-HG ranging from 1.5 – 5.5 µM (Fig. 3d). As expected, we did not observe a significant change in the dynamic range or the K_d_’ (Fig. 3e and Fig. 3f). In a background of higher concentrations of L-2-HG ranging from 15 – 55 µM, the dynamic range gradually decreased until plateauing at ∼1.3 at background L-2-HG concentrations ranging from 35 – 55 µM (Fig. 3e). Interestingly, higher concentrations of L-2-HG (15 – 55 µM) modestly increased the K_d_’ for D-2-HG by approximately 2 µM, demonstrating that an excess background and competition from L-2-HG only slightly inhibited D-2-HG binding (Fig 3f). In an attempt to improve the specificity of D2HGlo for D-2-HG over L-2-HG, we performed ligand docking studies of an AlphaFold generated model of DhdR to identify the potential binding site. The mutation of key residues in our predicted model did not affect D-2-HG or L-2-HG binding (Supplemental Fig. 4)^13^.

### Illumination of nuclear, cytoplasmic, and mitochondrial D-2-HG using D2HGlo

IDH1 is expressed in the cytosol and is primarily involved in producing NADPH for fatty acid biosynthesis, whereas IDH2 is present in the mitochondrial matrix and is a key player in the Krebs cycle. There is evidence that D-2-HG interferes with transcription in the nucleus and must ultimately be metabolized in the mitochondria by D-2-HG dehydrogenase^5,29–32^. Thus, D-2-HG may accumulate in the cytosol and multiple sub compartments of *IDH1* mutant cells to include the nucleus. To test the feasibility of using D2HGlo to interrogate cellular D-2-HG, we developed a cytosolic cellular probe, Cyto-D2HGlo, and assessed its response to exogenously applied D-2-HG and L-2-HG in HeLa cells. Cyto-D2HGlo responded to concentrations of D-2-HG ranging from 10 µM-10 mM, however, a significant difference in discrimination between D-2-HG and L-2-HG only occurred at a concentration of 100 µM (Supplemental Fig. 5).

Interestingly, we noticed that cells treated with concentrations of 2-HG ≥ 1 mM began to detach from the plate after a short time. To investigate if the apparent cytotoxic effect of D-2-HG and L-2-HG could be further characterized, we performed a cell viability assay using *IDH1*-R132H mutant U87MG, *IDH1* wildtype U87MG and SVG (normal human astrocyte) cells treated with D-2-HG or L-2-HG (Supplemental Fig. 6). We found concentrations at and above 1 mM of D-2-HG or L-2-HG to be cytotoxic.

Next, to assess whether endogenous D-2-HG could be detected in the cytoplasm, nucleus, and mitochondria of glioma cells, we developed two additional cellular probes, Nuc-D2HGlo and Mito-D2HGlo. Each exhibited proper localization in HeLa cells (Supplemental Fig. 7 and Supplemental Fig. 8) and U87MG cells (Fig. 4). Cyto-D2HGlo, Nuc-D2HGlo and Mito-D2HGlo were then expressed in *IDH1* wildtype and CRISPR-modified *IDH1*-R132H mutant U87MG cells. We found that the average FRET ratios of Cyto-D2HGlo, Nuc-D2HGlo, and Mito-D2HGlo were significantly elevated in the *IDH1* mutant cells compared to the *IDH1* wildtype cells (Fig. 4a-4c). Since the FRET ratio is a direct readout of D-2-HG concentration, these results suggest that *IDH1* mutations drive accumulation of D-2-HG in compartments other than the cytosol, including the nucleus where D-2-HG is known to have considerable epigenetic implications.

**Figure 4.**
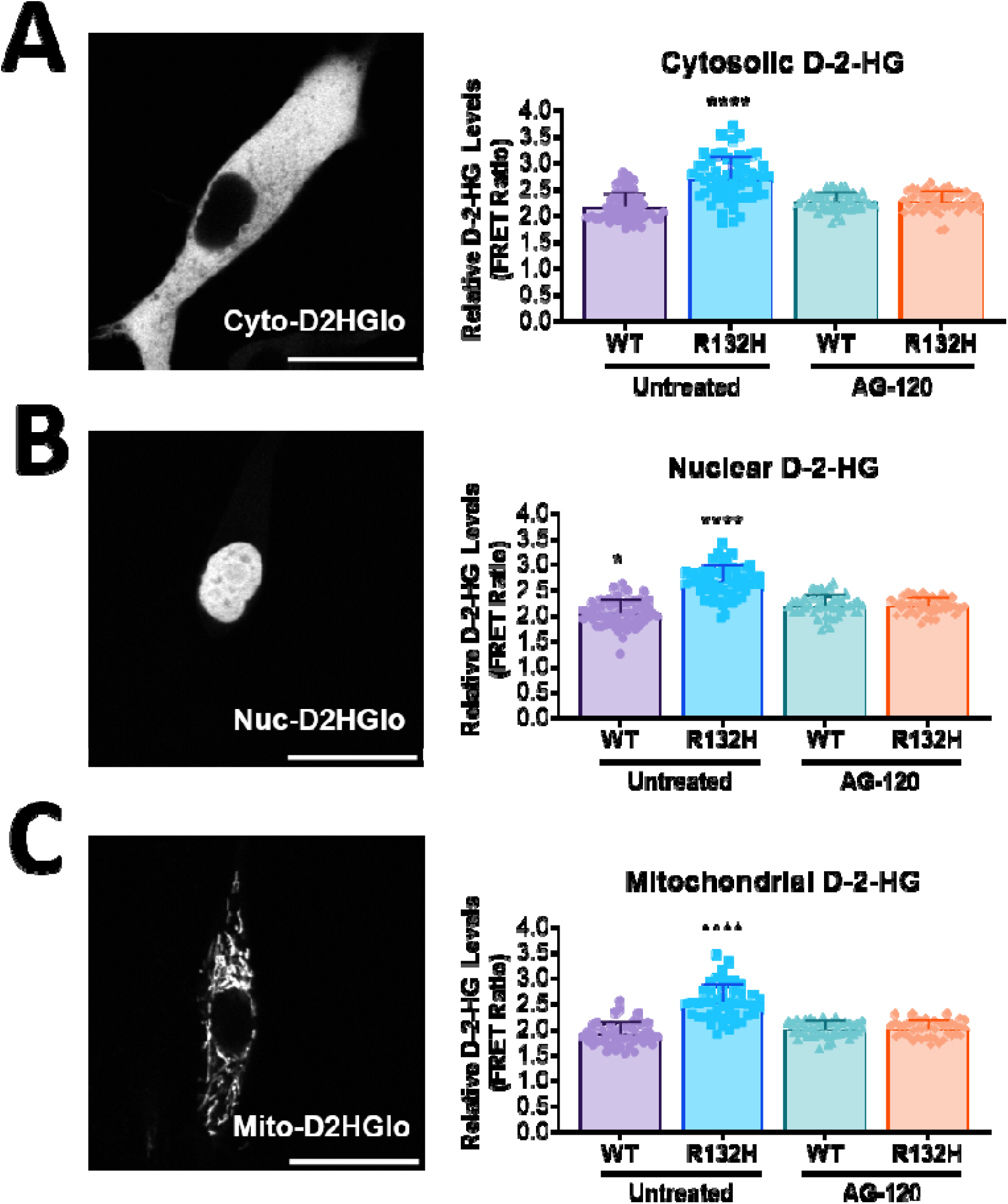
Subcellular targeting of the D2HGlo sensor reveals that D-2-HG levels are elevated in the cytosol, nucleus and mitochondria of *IDH1*-R132H mutant U87MG glioma cells. (A) *Left*: Representative image of a wildtype *IDH1* U87MG cell expressing Cyto-D2HGlo. Scale bar is 25 µm. *Right*: Comparison of the average FRET ratio in wildtype *IDH1* and *IDH1*-R132H mutant U87MG cells expressing Cyto-D2HGlo. The cells were grown in the presence or absence of the *IDH1*-R132H inhibitor AG-120 (3 µM) for 48 hours prior to collecting images. The average ± standard deviation is shown for n = 59 cells obtained from four independent experiments (untreated WT), n = 48 cells obtained from three independent experiments (AG-120-treated WT), n = 61 cells from four independent experiments (untreated R132H) and n = 47 cells from three independent experiments (AG-120-treated R132H). (B) *Left*: Representative image of a wildtype *IDH1* U87MG cell expressing Nuc-D2HGlo. Scale bar is 25 µm. *Right*: Comparison of the average FRET ratio in wildtype *IDH1* and *IDH1*-R132H mutant U87MG cells expressing Nuc-D2HGlo. Same treatments as panel A. The average ± standard deviation is shown for n = 59 cells obtained from four independent experiments (untreated WT), n = 45 cells obtained from three independent experiments (AG-120-treated WT), n = 58 cells from four independent experiments (untreated R132H) and n = 45 cells from three independent experiments (AG-120-treated R132H). (C) *Left*: Representative image of a wildtype *IDH1* U87MG cell expressing Mito-D2HGlo. Scale bar is 25 µm. *Right*: Comparison of the average FRET ratio in wildtype *IDH1* and *IDH1*-R132H mutant U87MG cells expressing Mito-D2HGlo. Same treatments as panel A. The average ± standard deviation is shown for n = 47 cells obtained from three independent experiments (untreated WT), n = 45 cells obtained from three independent experiments (AG-120-treated WT), n = 48 cells from three independent experiments (untreated R132H) and n = 45 cells from three independent experiments (AG-120-treated R132H). (A-D) Statistical analysis was performed using a one-way ANOVA test with *post hoc* Tukey (****, P < 0.0001 compared with untreated WT, AG-120-treated WT and AG-120-treated R132H; *, P < 0.05 compared with AG-120-treated WT and AG-120-treated R132H).

To validate the role of mutant *IDH1* in the response of D2HGlo to cellular D-2-HG, we utilized the FDA-approved mutant IDH1 inhibitor AG-120. Small molecule inhibitors of mutant IDH are a promising new targeted therapy effective in treating gliomas; however, their effect on D-2-HG levels in living cells has never been directly examined^33^. Real-time imaging of IDH1 mutant cells expressing Cyto-D2HGlo revealed that AG-120 did not alter the FRET ratio over a short time period (Supplemental Fig. 9). Therefore, we elected to treat *IDH1* wildtype and *IDH1*-R132H mutant U87MG cells with AG-120 for 48 hours prior to collecting FRET ratio images. We found that AG-120 (3 µM) reduced the average FRET ratio in all three compartments in *IDH1* mutant cells to levels comparable with *IDH1* wildtype cells (Fig. 4a). As expected, AG-120 had no effect on D-2-HG levels in wildtype *IDH1* U87MG cells. This evidence further supports the specificity of D2HGlo to endogenous D-2-HG *in-situ*, and these results are consistent with the findings of clinical trials involving both newly diagnosed and advanced cancer, which demonstrate a reduction of 2-HG to levels resembling those seen in healthy populations^34–36^.

While AG-120 was able to reduce the production of D-2-HG over a 48-hour period, no acute changes to the FRET ratio were observed. To determine whether real-time differences in cytosolic D-2-HG could be detected using Cyto-D2HGlo, we targeted other potential pathways upstream of mutant IDH-driven D-2-HG production. We utilized the glutaminase C inhibitor Compound 968 and 2-Deoxy-D-glucose (2-DG), an inhibitor of glycolysis. Acute application of 2-DG (10 mM) selectivity reduced the FRET ratio in *IDH1*-R132H mutant cells, whereas *IDH1* wildtype cells were unaffected (Supplemental Fig. 10b). In agreement with this finding, the concentration of D-2-HG in the supernatant of *IDH1*-R132H mutant cells that had been treated with 2-DG for 48 hours was reduced to levels comparable to untreated wildtype cells (Supplemental Fig. 10b). Acute application of Compound 968 (1 µM) did not elicit any changes in FRET ratio in wildtype or mutant cells expressing Cyto-D2HGlo (Supplemental Fig. 10c) nor did it effect D-2-HG accumulation in cell culture supernatants (Supplemental Figure 10d).

### D2HGlo detects D-2-HG in cell culture supernatant and human biological fluids

D-2-HG that is produced in cells is released into the extracellular environment^37^. Therefore, we examined whether D2HGlo could detect differences in D-2-HG concentrations in supernatants collected from wildtype *IDH1* and *IDH1*-R132H mutant U87MG cells. We found that the D-2-HG concentration of supernatant collected from wildtype cells was 6.56 ± 2.52 µM, while supernatant from *IDH1*-R132H cells contained 63.0 ± 2.7 µM (Fig. 5a). Treatment of *IDH1* mutant cells with AG-120 (3 µM) for 48 hours reduced the D-2-HG concentration to 2.79 ± 0.41 µM in *IDH1* wildtype cells while *IDH1*-R132H cells contained 2.88 ± 0.59 µM.

**Figure 5.**
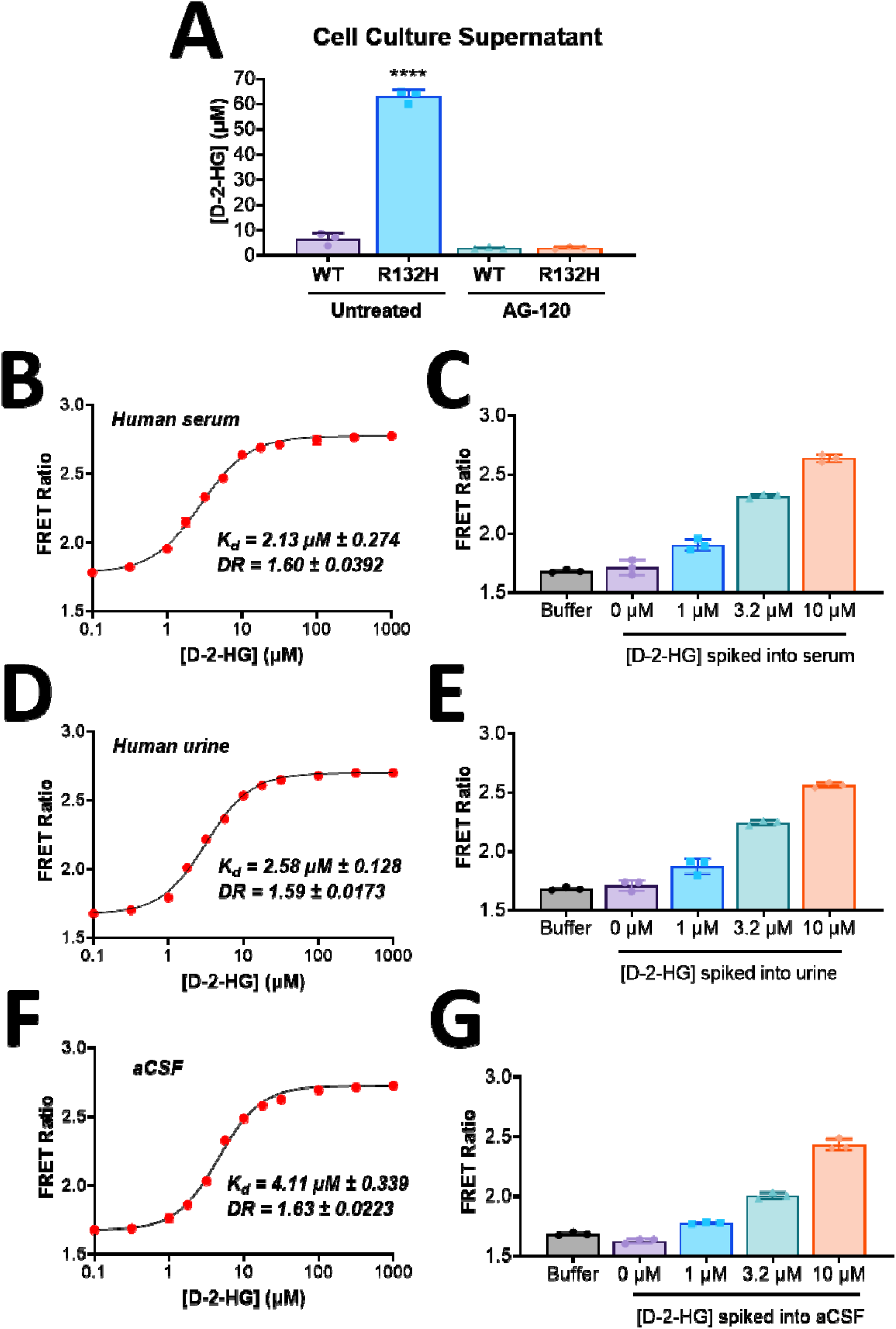
D2HGlo can be used as a diagnostic tool to monitor D-2-HG levels in human biological fluids. (A) Comparison of the D-2-HG concentration in supernatants collected from wildtype *IDH1* and *IDH1*-R132H mutant U87MG cells. Cells were grown in the presence or absence of the *IDH1*-R132H inhibitor AG-120 (3 µM) for 48 hours prior to collecting supernatant. (B) Titration of purified D2HGlo with increasing concentrations of D-2-HG (100 nM – 1 mM) spiked into human serum. (C) Average FRET ratio of D2HGlo in buffer, unspiked human serum and serum containing 1 µM, 3.2 µM or 10 µM D-2-HG. (D) Titration of purified D2HGlo with increasing concentrations of D-2-HG (100 nM – 1 mM) spiked into human urine. (E) Average FRET ratio of D2HGlo in buffer, unspiked human urine and urine containing 1 µM, 3.2 µM or 10 µM D-2-HG. (F) Titration of purified D2HGlo with increasing concentrations of D-2-HG (100 nM – 1 mM) spiked into aCSF. (G) Average FRET ratio of D2HGlo in buffer, unspiked aCSF and aCSF containing 1 µM, 3.2 µM or 10 µM D-2-HG. The K_d_’ and dynamic range for panels B, D and F are the result of three independent replicates.

The successful measurement of D-2-HG in culture medium suggested that D2HGlo may be amenable to detecting exogenous D-2-HG in biological fluids; thus, we next sought to quantify D-2-HG spiked into human serum, human urine, or artificial cerebrospinal fluid (aCSF) at concentrations ranging from 100 nM to 1 mM. The K_d_’ of D2HGlo for D-2-HG in human serum (Fig. 5b and Fig. 5c), human urine (Fig. 5d and Fig. 5e) and aCSF (Fig. 5f and Fig. 5g) was unaltered relative to measurements performed in buffer. Furthermore, the dynamic range showed little variation regardless of the biological matrix used; thus, the performance of D2HGlo is not diminished by serum, urine, or aCSF. These results support the potential use of D2HGlo to measure D-2-HG in clinical samples to detect IDH mutations and/or monitor patient therapy.

### D2HGlo successfully predicts the presence of *IDH1*-R132H mutations in glioma tumor samples

Following assessment of D2HGlo’s performance in biological fluids, we next sought to test the feasibility of measuring D-2-HG in the supernatant of archived tumor samples. Twenty glioma tumor samples with *IDH1* mutational status previously determined by immunohistochemistry (IHC) and loop-mediated isothermal amplification^38,39^ were assessed for the presence of D-2-HG. Substantially elevated FRET ratios were obtained for all *IDH1*-R132H tumor samples as compared to *IDH1* wildtype tumors (Fig. 6a). We found a highly significant increase in the mean FRET ratio upon analyzing *IDH1* mutant versus *IDH1* wildtype samples, with average FRET ratios of 2.05 ± 0.14 and 1.54 ± 0.05, respectively (Fig. 6b). Thus, D2HGlo successfully differentiated between *IDH1* wildtype and *IDH1* mutant samples with 100% concordance with the pathology report (Fig. 6c). D-2-HG concentrations in *IDH1*-R132H mutant tumor supernatants ranged between 11.1 and 72.3 µM and were not detectable in *IDH1*-R132 wildtype supernatants. These preliminary results suggest that a FRET ratio greater than 1.7, which correlates with a D-2-HG concentration of 1.87 µM, may be predictive of the presence of an *IDH1* mutation. This supports the feasibility of optimizing D2HGlo to intraoperatively diagnose the presence of an IDH mutation based on elevated D-2-HG levels.

**Figure 6.**
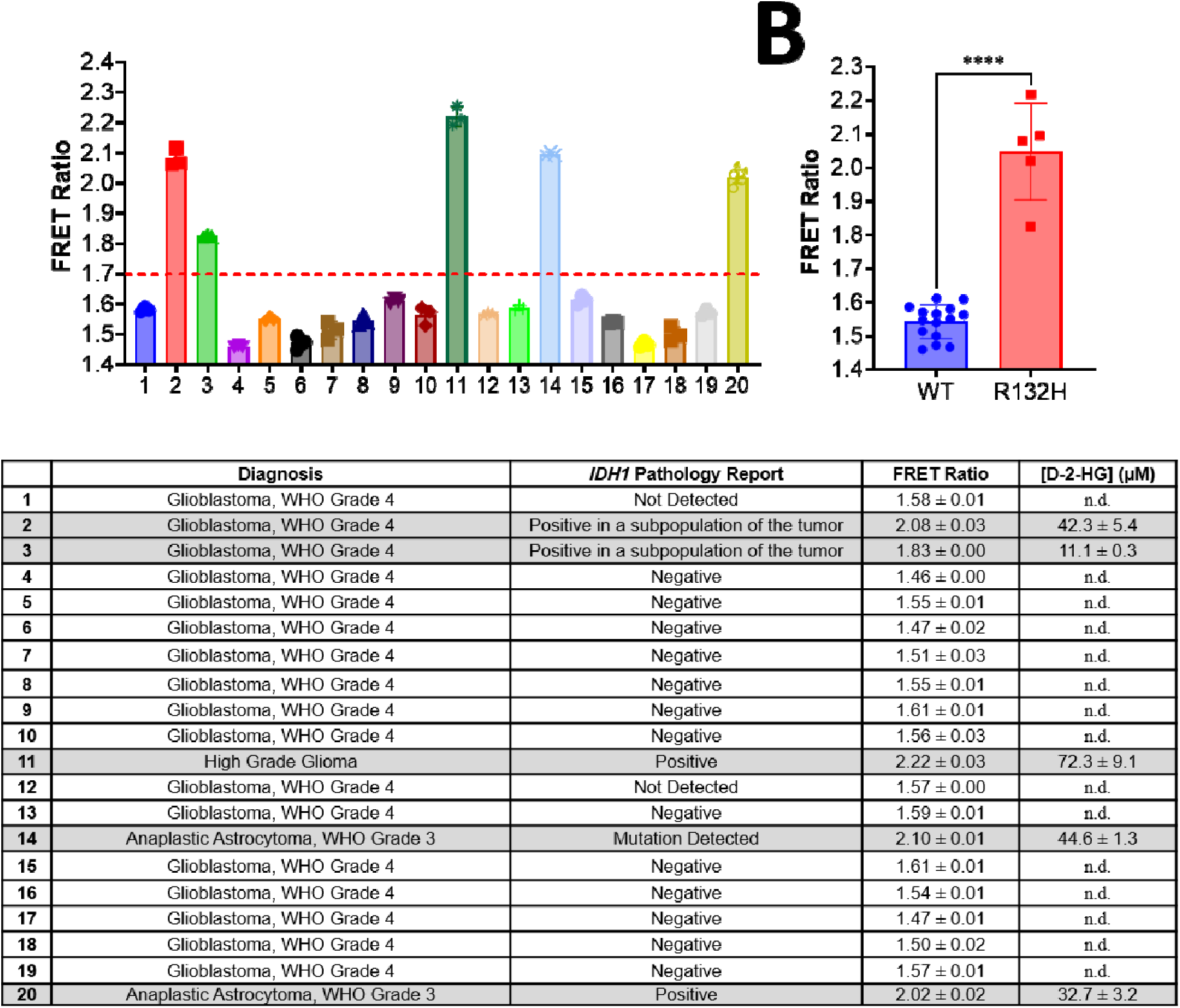
D2HGlo accurately predicts the *IDH1* mutational status of brain tumor samples from human patients. (A) The D2HGlo FRET ratio is shown for twenty brain tumor samples. 5 samples were derived from *IDH1*-R132H mutant individuals, and 15 were derived from wildtype *IDH1* samples. A FRET ratio of 1.7 was used as a threshold to predict IDH1 mutational status. (B) The average FRET ratio for wildtype *IDH1* compared with *IDH1* –R132H mutants. Statistical analysis was performed using an unpaired t-test (****, P < 0.0001). (C) Table showing diagnosis, pathology report, raw FRET ratio, and D-2-HG concentration for all twenty brain tumor samples. This pathology report was provided prior to the release of the fifth edition of the World Health Organization (WHO) Classification of Tumors of the Central Nervous System (CNS5). While samples 2 and 3 are listed as glioblastoma, new classifications place gliomas with IDH mutations in a separate category from GBM^40^.

## Discussion

Quantification of D-2-HG in samples has historically been challenging, and current assays are difficult, expensive, time-consuming, and/or do not function in a broad and biologically relevant range of concentrations. Discrimination between D-2-HG and L-2-HG is often ignored^41^ or difficult using commonly available techniques. We have developed a FRET-based sensor that can reliably measure D-2-HG in contrived clinical specimens and patient tumor supernatants.

Notably, D2HGlo does not respond to metabolites that are similar in structure to D-2-HG and can discriminate between D-2-HG and L-2-HG in the presence of physiologically relevant concentrations of L-2-HG. The sensitivity and specificity of D2HGlo demonstrates that it may be a powerful tool for quantifying D-2-HG via a liquid biopsy approach. Additionally, using targeted cellular probes of D2HGlo we detected elevated D-2-HG in several distinct subcellular compartments of *IDH1* mutant cells, including the nucleus.

Currently utilized techniques for the direct detection of IDH mutations such as genetic sequencing and IHC have higher cost and time requirements and are not amenable to remote testing. IHC is also limited to the detection of IDH1-R132H, whereas detection of elevated D-2-HG is inclusive of less common variants^42,43^. Quantification of D-2-HG can be performed using LC-MS^12,13,15,16^, ion-paired reverse-phase liquid chromatography^22^, GC-MS^18^, and MRS^17^. These methods do not easily discern between D-2-HG and L-2-HG, and performance time limits their utility. Commercially available fluorometric assays for detecting D-2-HG also exist but require a deproteinization step and 30-60 minute incubation. While currently available comparisons of D-2-HG levels in the body fluids of IDH wildtype and mutant patients vary based on technique and sample type, a positive correlation between mutational status and elevated D-2-HG has been demonstrated (Table 1)^1,10–14^.

There is an urgent need for a rapid and convenient liquid biopsy tool amenable for D-2-HG quantification in the clinical setting. Genetically encoded fluorescent sensors have been developed to quantify the concentration of metal ions and cellular metabolites *in situ*^44,45^, however, they are not routinely used as an *in vitro* diagnostic device. A FRET sensor capable of the specific detection of D-2-HG has recently been developed^46^ but is limited by a poor dynamic range and use at a physiologically impossible pH of 10. In comparison, D2HGlo performs at a physiologically relevant pH of 7.4 and quantifies D-2-HG within the currently documented disease range. In addition to its clinically relevant range of detection, D2HGlo offers robust quantification in complex matrices. D-2-HG may be measured within serum, plasma (EDTA plasma, sodium citrate plasma), urine, aCSF, cell culture media containing phenol red, and supernatants from glioma tumor lysates with no significant decrease in binding affinity or dynamic range in comparison to buffer. However, our results indicate that D2HGlo is not compatible with heparinized plasma or whole blood. Although we were not able to acquire patient serum, urine, or CSF samples, we demonstrate the feasibility of accurate quantification of D-2-HG using contrived samples in these matrices.

This assay could offer many advantages over current practices, including cost effectiveness and reducing patient distress. Results are available in approximately 15 minutes, and D-2-HG in biological fluids remains stable through several freeze-thaw cycles^11^ or exposure to excess heat^47^, making D2HGlo an attractive candidate for facilitating the remote diagnosis and disease monitoring of patients through mail in samples. Additionally, we have found D2HGlo’s performance to be highly reproducible. This is demonstrated by data in Figures 1b-c, 2b-d, 3b-f, 5a-g, and 6a-b, which are all representative of at least three independent sensor purifications.

While D2HGlo has a greater affinity for D-2-HG over L-2-HG, we recognize that its response to L-2-HG at higher concentrations may be a shortcoming in various potential applications. We found that at L-2-HG background concentrations below 10 µM, accurate quantification of D-2-HG reliably occurs; however, we found that accurate quantification of D-2-HG is increasingly hindered by elevated concentrations of L-2-HG (>10 µM) in a dose dependent manner. To address this, we attempted to improve enantiomer selectivity with site-directed mutagenesis of predicted binding domains but were unable to lower the affinity for D-2-HG or L-2-HG (Supplemental Fig. 4). While these results suggest we did not target the correct residues, it is highly probable that D-2-HG and L-2-HG share the same binding domain; thus, lowering the sensors affinity for L-2-HG will likely lower it for D-2-HG in tandem. Though D2HGlo’s utility in detecting *IDH1* mutations using serum, urine, and/or CSF must be assessed with clinical samples, successful discrimination between *IDH1* wildtype and *IDH1* mutant tumor samples was achieved in this study. L-2-HG did not appear to be present in appreciable amounts for any *IDH1* wildtype sample, as the FRET ratios obtained were not indicative of detectable amounts of ligand. Further, literature values of L-2-HG in the serum and CSF of humans without IDH mutations or metabolic disorders such as L-2-hydroxyglutaric aciduria are reported as ≤ 6.1 µM, suggesting D2HGlo is highly compatible with quantifying D-2-HG in these matrices.

In addition to the potential clinical applications of D2HGlo, we also demonstrate preliminary findings of its utility *in-situ*. We investigated the ability of D2HGlo to recognize subcellular differences in endogenous D-2-HG between *IDH1* mutant and *IDH1* wildtype cells by targeting the sensor to the nucleus, mitochondria, and cytoplasm. FRET ratios were significantly elevated in *IDH1* mutant cells in all three compartments, suggesting that D-2-HG accumulates throughout these cells. Our prediction that endogenous D-2-HG caused this increase in FRET ratio was supported by the inhibition of D-2-HG production through AG-120, a mutant IDH pharmaceutical inhibitor. 48-hour treatment with AG-120 caused the average FRET ratio for the nucleus, mitochondria, and cytoplasm of *IDH1* mutant cells to decrease to levels resembling those found in *IDH1* wildtype cells. Since no published tool currently exists to assess D-2-HG in living cells, these results offer findings of elevated D-2-HG within the nucleus, providing a potential mechanism for the abundance of epigenetic changes that are characteristic of IDH mutations. These findings also indicate that D2HGlo may be a valuable tool for further investigating the cellular distribution of D-2-HG and other cell-based studies of IDH mutant cells that have previously not been possible. However, as the crystal structure for DhdR has not been solved, we were unable to generate a dead sensor as a negative control in these experiments. Further studies such as the impact of pH and hypoxia on performance *in-situ* are warranted.

Collectively, our results suggest that D2HGlo may be used to infer the presence of IDH mutations preoperatively and/or monitor the efficacy of mutant IDH inhibitor therapy through liquid biopsies. The presence of *IDH1* mutations in glioma is associated with improved survival following aggressive surgical resection^48–50^, but no intraoperative methods are currently used. MRS may be employed to gauge total 2-HG levels preoperatively, but this technique is limited by high cost, operator dependence, and low signal-to-noise ratio which require lesions to be at least several millimeters in volume and sufficiently distant from fluid-brain or air-fluid interfaces. As D2HGlo could accurately predict *IDH1* mutational status in glioma tumor supernatants, the intraoperative use of D2HGlo may prove useful if altering the extent of resection based on IDH mutational status is desired. For future clinical implementation, we intend to validate D2HGlo in accordance with CLIA standards for its use as a diagnostic and predictive tool and envision the development of a lyophilized sensor that can be resuspended using sample buffer, water, or patient samples. In addition to clinical applications, we also present D2HGlo cellular probes which facilitated the study of D-2-HG distribution *in-situ*. These probes may play a valuable role in further characterizing the mechanism of D-2-HG driven oncogenesis in IDH mutant cells.^48–51^

## Materials and Methods

### Chemicals and reagents

Oxaloacetic acid, α-ketoglutaric acid, succinic acid, L-glutamic acid, sodium hydrogen DL-malate, sodium L-lactate, and 2-Deoxy-D-glucose were purchased from Thermo Fisher Scientific (Waltham, MA). Sodium citrate was purchased from Ward’s Science. Fumarate was purchased from TCI Chemical Company. Sodium pyruvate was purchased from Beantown Chemical. D-2-HG, L-2-HG, isocitrate, AG-120, 2R-Octyl-11-hydroxyglutarate, 2S-Octyl-11-hydroxyglutarate, and glutaminase inhibitor (compound 968) were purchased from Cayman Chemical (Ann Arbor, MI).

### Molecular Cloning

The gene encoding the transcription factor DhdR from *Achromobacter denitrificans* NBRC 15125 (ADE01S_RS31870) was purchased as a gene fragment from Azenta Life Sciences (Burlington, MA). The DhdR gene was PCR amplified using primers containing SphI and SacI restriction sites and inserted into a pBAD vector containing the fluorescent proteins ECFP and cpVenus173 (Supplemental Table 2). Both fluorescent proteins originated from the plasmid pcDNA3.1(+)-NES-ZapCV2 (cpV143) (Addgene Plasmid #36231). Three variants of the parent DhdR-based FRET sensor were generated by removing the full-length DhdR domain and inserting PCR amplified truncated versions of DhdR in its place. For mammalian expression of D2HGlo, D2HGlo was PCR amplified from the pBAD vector and transferred to one of two pcDNA3 backbones. To generate Cyto-D2HGlo and Nuc-D2HGlo, pcDNA3.1(+)-NES-ZapCV2 (cpV143) (Addgene Plasmid #36231) was used as a destination vector. Both D2HGlo and the vector were digested with BamHI and EcoRI (Cyto-D2HGlo) or KpnI and EcoRI (Nuc-DH2Glo). Cyto-D2HGlo contained the nuclear export signal (NES) MLQLPPLERLTL at the N-terminus, and Nuc-D2HGlo included the nuclear localization signal (NLS) MPKKKRKVEDA at the N-terminus. To generate Mito-D2HGlo, pcDNA-mito-ZapCY (Addgene Plasmid #36231) was used as the destination vector. The restriction enzymes BamHI and EcoRI were used for restriction cloning. This resulted in a mitochondrial D2HGlo construct with four copies of CoxVII (YVRPDAAAAAGLDRLGPAAPSAARQDPFVG) at the N-terminus. All DNA constructs were transformed into Invitrogen™ Subcloning Efficiency™ DH5α Competent Cells, and colonies were selected and cultured in LB broth containing 100 ug/mL ampicillin. Plasmid DNA was purified using E.Z.N.A Plasmid DNA Mini Kit I (Omega Bio-Tek, D6948). All DNA constructs were confirmed by Sanger sequencing.

### Site-Directed Mutagenesis

AlphaFold was utilized to convert the primary sequence of DhdR into a 3D structure, then SwissDock was used to identify the most stable binding mode for DhdR with D-2-HG or L-2-HG ligands. SwissDock identified that L-2-HG formed an interaction with T52 and D-2-HG formed an interaction with S69. PCR was then utilized to generate three constructs containing mutations anticipated to impact 2-HG binding (T52A, S69A, and S69T). Mutations were produced by incorporating the desired base pair changes into the primer sequence, amplifying the D2HGlo plasmid, and digesting the original construct with DPNI. Prepared constructs were transformed into competent bacteria followed by protein purification and sensor assessment via titrations of D-2-HG and L-2-HG as previously described. Primer sequences can be found in Supplemental Table 2.

### Protein Expression and Purification

The DhdR-based FRET construct encoded by the pBAD plasmid were transformed into One Shot TOP10 Chemically Competent *E. coli* (Thermo Fisher Scientific, Waltham, MA). Colonies were selected and grown up at 37°C in 2xYT microbial growth media containing 100 µg/mL ampicillin. Once the culture reached an OD_600_ of approximately 0.6, protein expression was induced with the addition of 0.2% L-arabinose. Cultures were grown at 22°C for 24 hours. Bacterial cell lysis was performed using B-PER reagent (Thermo Fisher Scientific) supplemented with lysozyme (50 mg/mL) and DNase I (2,500 U/mL). The resulting crude lysate was cleared by centrifugation, and the supernatant was sterile filtered. D2HGlo was purified using Cobalt Metal Affinity Resin (Takara Bio). The resin was washed with equilibration buffer (50 mM Tris, 300 mM NaCl, pH 7.4) prior to the addition of the cleared lysate. The column was rinsed with wash buffer (50 mM Tris, 300 mM NaCl, 20 mM imidazole) twice before addition of the elution buffer (50 mM Tris, 300 mM NaCl, 250 mM imidazole). Using a PD-10 desalting column, the eluent was exchanged into an experimental buffer containing 150 mM HEPES, 100 mM NaCl and 10% glycerol (pH 7.4). To examine the influence of pH on D2HGlo function, the experimental buffer was buffered to pH 6.5 or pH 8. In all experiments, the concentration of purified FRET sensor was determined based on absorbance of the acceptor fluorescent protein cpVenus173 (ε = 92,000 M^-1^cm^-1^ at 515 nm).

### *In vitro* characterization of purified D2HGlo

Purified D2HGlo was diluted to 5 µM in experimental buffer for all *in vitro* fluorescence measurements. D-2-HG was prepared as a 50 mM stock in phosphate buffered saline (PBS) at pH 7.4. Serial dilutions were performed to achieve final concentrations of D-2-HG between 10 nM – 1 mM. L-2-HG and other cellular metabolites were prepared in the same manner. In a black-walled clear bottom 96-well plate, 90 µL of purified sensor protein was mixed with 10 µL of a 10X stock concentration. Fluorescence measurements were performed using a Synergy Multi-Mode Plate Reader (BioTek Instruments, Winooski, VT). For the collection of emission spectra, the imaging parameters were: 420/10 nm excitation and 450-650 nm emission window (1 nm stepsize). For the D-2-HG binding studies, the imaging parameters were: 440 nm excitation/485 nm emission and 440 nm excitation/531 emission. The FRET ratio was calculated by dividing the fluorescent signal collected at 531 nm by the fluorescent signal collected at 485 nm. The FRET ratio was plotted against the log_10_ concentration of D-2-HG. The data points were fit with the following equation to obtain a K_d_’: FRET Ratio = FRET_bound_ x [D-2-HG]^n^ + FRET_unbound_ x K ^n^) / (CFP x [D-2-HG]^n^ + CFP x K ^n^, where FRET is the maximum FRET ratio and FRET_unbound_ is the minimum FRET ratio^52^. For temperature-based characterization, varying concentrations of D-2-HG were combined with 5 <u>µM</u> sensor, incubated at 30°C or 37°C for 15 minutes, then fluorescence measurements were obtained immediately. For pH experiments, sensor was prepared in titration buffer with a final pH of 6.5, 7.5, or 8.5 then analyzed for D-2-HG binding capacity as previously specified.

### Cell Culture and Transfection

*IDH1* wildtype U87MG, CRISPR edited *IDH1*-R132H Isogenic U87HTB-141G™ (ATCC, Manassas, VA), and HeLa cells (ATCC, Manassas, VA) were cultured in DMEM (Lonza, Portsmouth, NH) supplemented with 10% fetal bovine serum (Atlanta Biologicals, Norcross, GA) and 1% penicillin/streptomycin/amphotericin B (PSA) (Lonza, Portsmouth, NH,) at 37°C and 5% CO_2_. For imaging experiments, cells were plated in 6-well dishes at 70-80% confluence. Cells were transfected 24 hours after plating with 1.0 µg D2HGlo constructs and 3.0 µL Trans-IT®-LT1 transfection reagent (Mirus Bio, Madison, WI). The cells were allowed to express for 48 hours before fluorescence imaging.

### Fluorescence Live-Cell Imaging

IDH1 wildtype U87MG, CRISPR edited IDH1-R132H Isogenic U87HTB-141G™ cells and HeLa cells were imaged in 8-chamber coverslips (Ibidi, Fitchburg, WI). The media was exchanged for HEPES-buffered Hank’s Balanced Salt Solution (140 mM NaCl, 5 mM KCl, 1 mM CaCl_2_, 4 mM MgSO_4_, 5 mM MgCl_2_, 3 mM Na_2_PO4, 4 mM PO_4_, 6 mM glucose, 4 mM NaHCO_3_, pH 7.4). Cells expressing the D2HGlo sensors were imaged using an Olympus FLUOVIEW FV3000 Confocal Microscope. The D2HGlo sensors were excited at 458 nm and the fluorescent signal was collected using a BA 480-495 emission filter (ECFP) and a BA 535-565 emission filter (FRET). The FRET ratio was determined by dividing the FRET signal by the ECFP signal. For real-time FRET experiments of HeLa cells expressing Cyto-D2HGlo, a baseline FRET ratio was collected for one minute prior to addition of D-2-HG or L-2-HG at 0.01 mM, 0.1 mM, 1 mM or 10 mM. The FRET ratio was collected until a maximum FRET ratio had been achieved or the cells had died. The maximum FRET ratio was determined for the entire time-course and normalized to the baseline FRET ratio. For long-term FRET experiments, ratiometric FRET images of U87MG or CRISPR edited IDH1-R132H Isogenic U87HTB-141G™ cells that had been transfected with Cyto-D2HGlo, Nuc-D2HGlo or Mito-D2HGlo were collected. A region of interest was drawn on each cell, and the average FRET ratio within each ROI was determined. In some sets of experiments, cells were treated with AG-120 (3 µM) for 48 hours prior to FRET ratio collection. Alternatively, glutaminase inhibitor (1 µM), or 2-DG (10 mM) were added to imaging wells after collecting baseline FRET values for 30 seconds.

### Cell Viability Assays

U87MG, *IDH1*-R132H mutant U87MG, and SVG (ATCC, Manassas, VA, CRL-8621) cells were plated at 10,000 cells per well in an opaque 96-well plate with complete media and incubation conditions as described previously. Cells were incubated overnight, then media was then replaced with DMEM containing 10% FBS, 1% PSA, and a concentration of cell permeable D-2-HG (2R-Octyl-11-hydroxyglutarate), or L-2-HG (2S-Octyl-11 hydroxyglutarate) ranging from 4 mM-0 µM. Cells were incubated for 48 hours, then assessed for viability with CellTiter-Glo® (Promega) according to protocol. Luminescence was measured on a Synergy Multi-Mode Plate Reader.

### Cell Culture Supernatant

U87MG *IDH1* wildtype and mutant cells were plated at 60% confluence in 6-well plates. For some experiments, AG-120 in DMSO (3 µM), Glutaminase inhibitor (1 µM), or 2-DG (10 mM) was added to the cell culture media before plating the cells. For all conditions and cell lines, the cell culture supernatant was collected after 48 hours of incubation at 37°C. 10 µL of cell culture media was mixed with 90 µL of purified D2HGlo (5 µM), and the FRET ratios were collected. D-2-HG levels were estimated by using the titration curve as a calibration curve and converting the FRET ratios of unknowns into concentrations.

### Biological Fluids

Urine and serum were collected from three healthy donors while aCSF was purchased (Tocris Bioscience, Bristol, England). Biological fluids were spiked with D-2-HG at concentrations ranging from 10 nM to 1 mM. Fluorescence was collected at 485 nm and 531 nm, and the FRET ratio was determined. Binding constants were determined as previously described.

### Tumor Lysates

Advocate Aurora Research Institute, LLC, Milwaukee, WI, generously provided Archived Tumor Samples with known *IDH1* mutational status (Aurora IRB# 14–79), which was further confirmed using CPNA-LAMP. Tumor lysates were created by homogenizing approximately 0.1 gram of tumor in 500 µL of physiological saline. A portion of the archived samples were identified as post-processing cells (samples 10, 12, 15, 16, and 18) and were analyzed directly. Prior to measurement with FRET, the samples were thawed and centrifuged at 16,000XG for 4 minutes.

### Statistics and Reproducibility

At least three independent experiments were performed for each assay. Data is shown as either representative experiments (binding curves) or individual data points (bar graphs). Statistical analysis was performed using GraphPad Prism 10.1.2.

### Funding

This work was supported by the Upper Michigan Brain Tumor Center and the CHSPS grant (College of Health Sciences and Professional Studies) at Northern Michigan University.

### Author Contributions

K.A.C and W.W.L.K conducted all experiments and performed data analysis. P.B.M and M.J.J contributed to the acquisition of funding, experimental design, and the acquisition of clinical samples. R.J.W contributed to the acquisition of funding. D.O.K assembled data found in table 1 and offered insight toward the clinical applications of D2HGlo. E.P.S.P contributed to the conceptualization and design of all sensor constructs, experimental design, preparation of figures, and acquisition of funding. K.A.C and E.P.S.P prepared the original manuscript, and all authors contributed to its editing and review.

### Competing interests

The authors declare no competing interests.

### Data and materials availability

All data needed to evaluate the conclusions in the paper are present in the paper or the supplementary materials.

## Notes

### Competing Interest Statement

The authors have declared no competing interest.

## References

1 Dang, L. et al. Cancer-associated *IDH1* mutations produce 2-hydroxyglutarate. Nature 462, 739–744 (2009). 10.1038/nature08617

2 Raimondi, V., Ciotti, G., Gottardi, M. & Ciccarese, F. 2-Hydroxyglutarate in acute myeloid leukemia: A Journey from Pathogenesis to Therapies. Biomedicines 10 (2022). 10.3390/biomedicines10061359

3 Philip, B. et al. Mutant *IDH1* promotes glioma formation *in vivo*. Cell Rep 23, 1553–1564 (2018). 10.1016/j.celrep.2018.03.133

4 Ye, D., Guan, K. L. & Xiong, Y. Metabolism, activity, and targeting of D-and L-2-Hydroxyglutarates. Trends Cancer 4, 151–165 (2018). 10.1016/j.trecan.2017.12.005

5 Turcan, S., et al. *IDH1* mutation is sufficient to establish the glioma hypermethylator phenotype. Nature 483, 479–483 (2012). 10.1038/nature10866

6 Malta, T. M. et al. Glioma CpG island methylator phenotype (G-CIMP): biological and clinical implications. Neuro Oncol 20, 608–620 (2018). 10.1093/neuonc/nox183

7 Cui, D. et al. R132H mutation in *IDH1* gene reduces proliferation, cell survival and invasion of human glioma by downregulating Wnt/β-catenin signaling. Int J Biochem Cell Biol 73, 72–81 (2016). 10.1016/j.biocel.2016.02.007

8 Hartmann, C. et al. Patients with IDH1 wild type anaplastic astrocytomas exhibit worse prognosis than *IDH1*-mutated glioblastomas, and IDH1 mutation status accounts for the unfavorable prognostic effect of higher age: implications for classification of gliomas. Acta Neuropathol 120, 707–718 (2010). 10.1007/s00401-010-0781-z

9 Delahousse, J. et al. Circulating oncometabolite D-2-hydroxyglutarate enantiomer is a surrogate marker of isocitrate dehydrogenase–mutated intrahepatic cholangiocarcinomas. European Journal of Cancer 90, 83–91 (2018). 10.1016/j.ejca.2017.11.024

10 Lee, C. L. et al. Circulating oncometabolite 2-hydroxyglutarate as a potential biomarker for isocitrate dehydrogenase (*IDH1/2*) mutant cholangiocarcinoma. Mol Cancer Ther 23, 394–399 (2024). 10.1158/1535-7163.Mct-23-0460

11 Kalinina, J. et al. Selective detection of the D-enantiomer of 2-Hydroxyglutarate in the CSF of glioma patients with mutated isocitrate dehydrogenase. Clin Cancer Res 22, 6256–6265 (2016). 10.1158/1078-0432.CCR-15-2965

12 Fujita, Y., et al. IDH1 p.R132H ctDNA and D-2-hydroxyglutarate as CSF biomarkers in patients with IDH-mutant gliomas. J Neurooncol 159, 261–270 (2022). 10.1007/s11060-022-04060-1

13 Tuna, G. et al. Minimally invasive detection of IDH1 mutation with cell-free circulating tumor DNA and D-2-Hydroxyglutarate, D/L-2-Hydroxyglutarate ratio in gliomas. J Neuropathol Exp Neurol 81, 502–510 (2022). 10.1093/jnen/nlac036

14 Lombardi, G. et al. Diagnostic value of plasma and urinary 2-hydroxyglutarate to identify patients with isocitrate dehydrogenase-mutated glioma. Oncologist 20, 562–567 (2015). 10.1634/theoncologist.2014-0266

15 Struys, E. et al. Measurement of urinary D-and L-2-hydroxyglutarate enantiomers by stable-isotope-dilution liquid chromatography-tandem mass spectrometry after derivatization with diacetyl-L-tartaric anhydride. Clin Chem 50, 1391–1395 (2004). 10.1373/clinchem.2004.033399

16 Zhang, C. et al. Serum D-2-hydroxyglutarate and the ratio of D-2HG/L-2HG predict *IDH* mutation in acute myeloid leukemia. EJHaem 4, 723–727 (2023). 10.1002/jha2.723

17 Choi, C. et al. 2-hydroxyglutarate detection by magnetic resonance spectroscopy in *IDH*-mutated patients with gliomas. Nat Med 18, 624–629 (2012). 10.1038/nm.2682

18 Fernandez-Galan, E. et al. Validation of a routine gas chromatography mass spectrometry method for 2-hydroxyglutarate quantification in human serum as a screening tool for detection of idh mutations. J Chromatogr B Analyt Technol Biomed Life Sci 1083, 28–34 (2018). 10.1016/j.jchromb.2018.02.038

19 Mellinghoff, I. K. et al. Vorasidenib in *IDH1*-or *IDH2*-mutant low-grade glioma. N Engl J Med 389, 589–601 (2023). 10.1056/NEJMoa2304194

20 Strain, S. K., et al. Enantioseparation and detection of (R)-2-Hydroxyglutarate and (S)-2-Hydroxyglutarate by chiral gas chromatography–triple quadrupole mass spectrometry. Metabolomics, 89–100 (2021). 10.1007/978-1-0716-0864-7_8

21 Sim, H. W. et al. Tissue 2-Hydroxyglutarate as a biomarker for isocitrate dehydrogenase mutations in gliomas. Clin Cancer Res 25, 3366–3373 (2019). 10.1158/1078-0432.Ccr-18-3205

22 Fathi, A. T. et al. Elevation of urinary 2-Hydroxyglutarate in IDH-Mutant glioma. Oncologist 21, 214–219 (2016). 10.1634/theoncologist.2015-0342

23 Capper, D. et al. 2-Hydroxyglutarate concentration in serum from patients with gliomas does not correlate with *IDH1/2* mutation status or tumor size. Int J Cancer 131, 766–768 (2012). 10.1002/ijc.26425

24 Xiao, D. et al. A D-2-hydroxyglutarate biosensor based on specific transcriptional regulator DhdR. Nat Commun 12, 7108 (2021). 10.1038/s41467-021-27357-7

25 Ewald, J. C. et al. Engineering genetically encoded nanosensors for real-time *in vivo* measurements of citrate concentrations. PLoS One 6, e28245 (2011). 10.1371/journal.pone.0028245

26 Luddecke, J. et al. P(II) Protein-derived FRET sensors for quantification and live-cell imaging of 2-Oxoglutarate. Sci Rep 7, 1437 (2017). 10.1038/s41598-017-01440-w

27 Rzem, R., et al. L-2-hydroxyglutaric aciduria, a defect of metabolite repair. J Inherit Metab Dis 30, 681–689 (2007). 10.1007/s10545-007-0487-0

28 Intlekofer, A. M. et al. L-2-Hydroxyglutarate production arises from noncanonical enzyme function at acidic pH. Nat Chem Biol 13, 494–500 (2017). 10.1038/nchembio.2307

29 Chowdhury, R. et al. The oncometabolite 2-hydroxyglutarate inhibits histone lysine demethylases. EMBO Rep 12, 463–469 (2011). 10.1038/embor.2011.43

30 Kusi, M. et al. 2-Hydroxyglutarate destabilizes chromatin regulatory landscape and lineage fidelity to promote cellular heterogeneity. Cell Rep 38, 110220 (2022). 10.1016/j.celrep.2021.110220

31 Yang, Z. et al. 2-HG inhibits necroptosis by stimulating *DNMT1*-dependent hypermethylation of the *RIP3* promoter. Cell Rep 19, 1846–1857 (2017). 10.1016/j.celrep.2017.05.012

32 Ceccarelli, M. et al. Molecular profiling reveals biologically discrete subsets and pathways of progression in diffuse glioma. Cell 164, 550–563 (2016). 10.1016/j.cell.2015.12.028

33 Popovici-Muller, J. et al. Discovery of AG-120 (Ivosidenib): A first-in-class mutant IDH1 inhibitor for the treatment of *IDH1* mutant cancers. ACS Med Chem Lett 9, 300–305 (2018). 10.1021/acsmedchemlett.7b00421

34 Fan, B. et al. Pharmacokinetic/pharmacodynamic evaluation of Ivosidenib or Enasidenib combined with intensive induction and consolidation chemotherapy in patients with newly diagnosed *IDH1/2*-mutant acute myeloid leukemia. Clin Pharmacol Drug Dev 11, 429–441 (2022). 10.1002/cpdd.1067

35 Fan, B. et al. Clinical pharmacokinetics and pharmacodynamics of ivosidenib in patients with advanced hematologic malignancies with an *IDH1* mutation. Cancer Chemother Pharmacol 85, 959–968 (2020). 10.1007/s00280-020-04064-6

36 Fan, B. et al. Clinical pharmacokinetics and pharmacodynamics of ivosidenib, an oral, targeted inhibitor of mutant *IDH1*, in patients with advanced solid tumors. Invest New Drugs 38, 433–444 (2020). 10.1007/s10637-019-00771-x

37 Bunse, L. et al. Suppression of antitumor T cell immunity by the oncometabolite (R)-2-hydroxyglutarate. Nature Medicine 24, 1192–1203 (2018). 10.1038/s41591-018-0095-6

38 Choate, K. A. et al. Rapid extraction-free detection of the R132H isocitrate dehydrogenase mutation in glioma using colorimetric peptide nucleic acid-loop mediated isothermal amplification (CPNA-LAMP). PLoS One 18, e0291666 (2023). 10.1371/journal.pone.0291666

39 Choate, K. A. et al. Rapid *IDH1*-R132 genotyping panel utilizing locked nucleic acid loop-mediated isothermal amplification (LNA-LAMP). Biology Methods and Protocols (2024). 10.1093/biomethods/bpae012

40 Louis, D. N. et al. The 2021 WHO classification of tumors of the central nervous system: a summary. Neuro Oncol 23, 1231–1251 (2021). 10.1093/neuonc/noab106

41 Wang, J. H. et al. Prognostic significance of 2-hydroxyglutarate levels in acute myeloid leukemia in China. Proc Natl Acad Sci U S A 110, 17017–17022 (2013). 10.1073/pnas.1315558110

42 Nakagawa, M. et al. Clinical usefulness of 2-hydroxyglutarate as a biomarker in IDH-mutant chondrosarcoma. J Bone Oncol 34, 100430 (2022). 10.1016/j.jbo.2022.100430

43 Pusch, S. et al. D-2-Hydroxyglutarate producing neo-enzymatic activity inversely correlates with frequency of the type of isocitrate dehydrogenase 1 mutations found in glioma. Acta Neuropathol Commun 2, 19 (2014). 10.1186/2051-5960-2-19

44 Pratt, E. P. S. et al. Regulation of cAMP accumulation and activity by distinct phosphodiesterase subtypes in INS-1 cells and human pancreatic β-cells. PLoS One 14, e0215188 (2019). 10.1371/journal.pone.0215188

45 Pratt, E. P. S. et al. Tools and techniques for illuminating the cell biology of zinc. Biochim Biophys Acta Mol Cell Res 1868, 118865 (2021). 10.1016/j.bbamcr.2020.118865

46 Xiao, D. et al. A Förster resonance energy transfer-based D-2-hydroxyglutarate biosensor. Sensors and Actuators B: Chemical 385, 133681 (2023). 10.1016/j.snb.2023.133681

47 Voelxen, N. F. et al. Quantitative imaging of D-2-Hydroxyglutarate in selected histological tissue areas by a novel bioluminescence technique. Front Oncol 6, 46 (2016). 10.3389/fonc.2016.00046

48 Beiko, J., et al. *IDH1* mutant malignant astrocytomas are more amenable to surgical resection and have a survival benefit associated with maximal surgical resection. Neuro Oncol 16, 81–91 (2014). 10.1093/neuonc/not159

49 Cahill, D. P. Extent of resection of glioblastoma: A critical evaluation in the molecular era. Neurosurg Clin N Am 32, 23–29 (2021). 10.1016/j.nec.2020.09.006

50 Jakola, A. S. et al. The impact of resection in IDH-mutant WHO grade 2 gliomas: a retrospective population-based parallel cohort study. J Neurosurg, 1–8 (2022). 10.3171/2022.1.JNS212514

51 Molinaro, A. M. et al. Association of maximal extent of resection of contrast-enhanced and non-contrast-enhanced tumor with survival within molecular subgroups of patients with newly diagnosed glioblastoma. JAMA Oncol 6, 495–503 (2020). 10.1001/jamaoncol.2019.6143

52 Pomorski, A. et al. Method for accurate determination of dissociation constants of optical ratiometric systems: chemical probes, genetically encoded sensors, and interacting molecules. Anal Chem 85, 11479–11486 (2013). 10.1021/ac402637h

