## Supplemental figures for "A genetically encoded fluorescent sensor enables sensitive and specific detection of IDH mutant associated oncometabolite D-2-hydroxyglutarate"

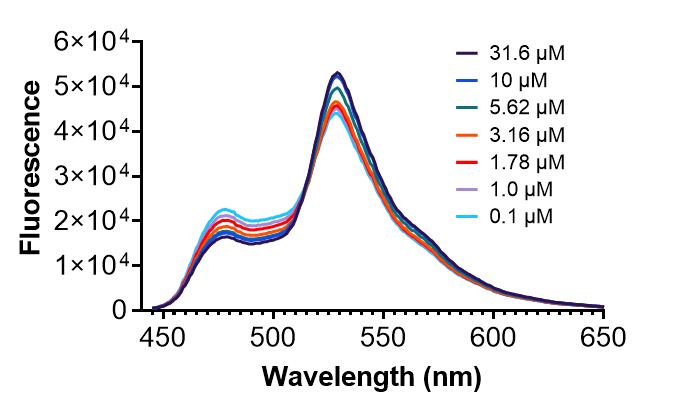

**Supplemental Figure 1. Emission spectrum of purified D2HGlo.** The purified sensor was exposed to D-2-HG at concentrations ranging from 0.1-31.6 µM. The FRET sensor was excited at 440 nm, and the emission spectrum window was collected between 450-650 nm. As the concentration of D-2-HG increased, the 482 nm peak corresponding with ECFP increased and the 531 nm peak increased. Thus, this supports a model in which D-2-HG binding DhdR results in increased FRET between the two fluorescent proteins.

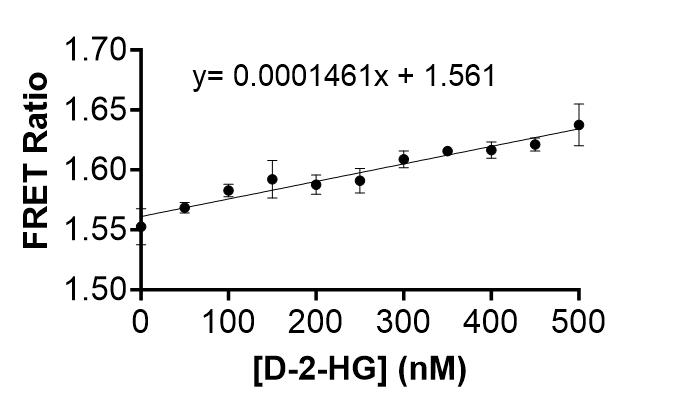

**Supplemental Figure 2. Limit of detection of D2HGlo.** Purified D2HGlo was exposed to concentrations of D-2-HG, ranging from 50 nM to 500 nM, and the FRET ratio was collected at each concentration.The limit of detection (LOD) was determined using the following equation: LOD = 3ơ/S, where ơ is the standard deviation of the FRET ratio measurements collected from the blank (no D-2-HG present) and S is the slope obtained from linear regression analysis. Based on this calculation, the LOD of D2HGlo was determined to be 308 nM.

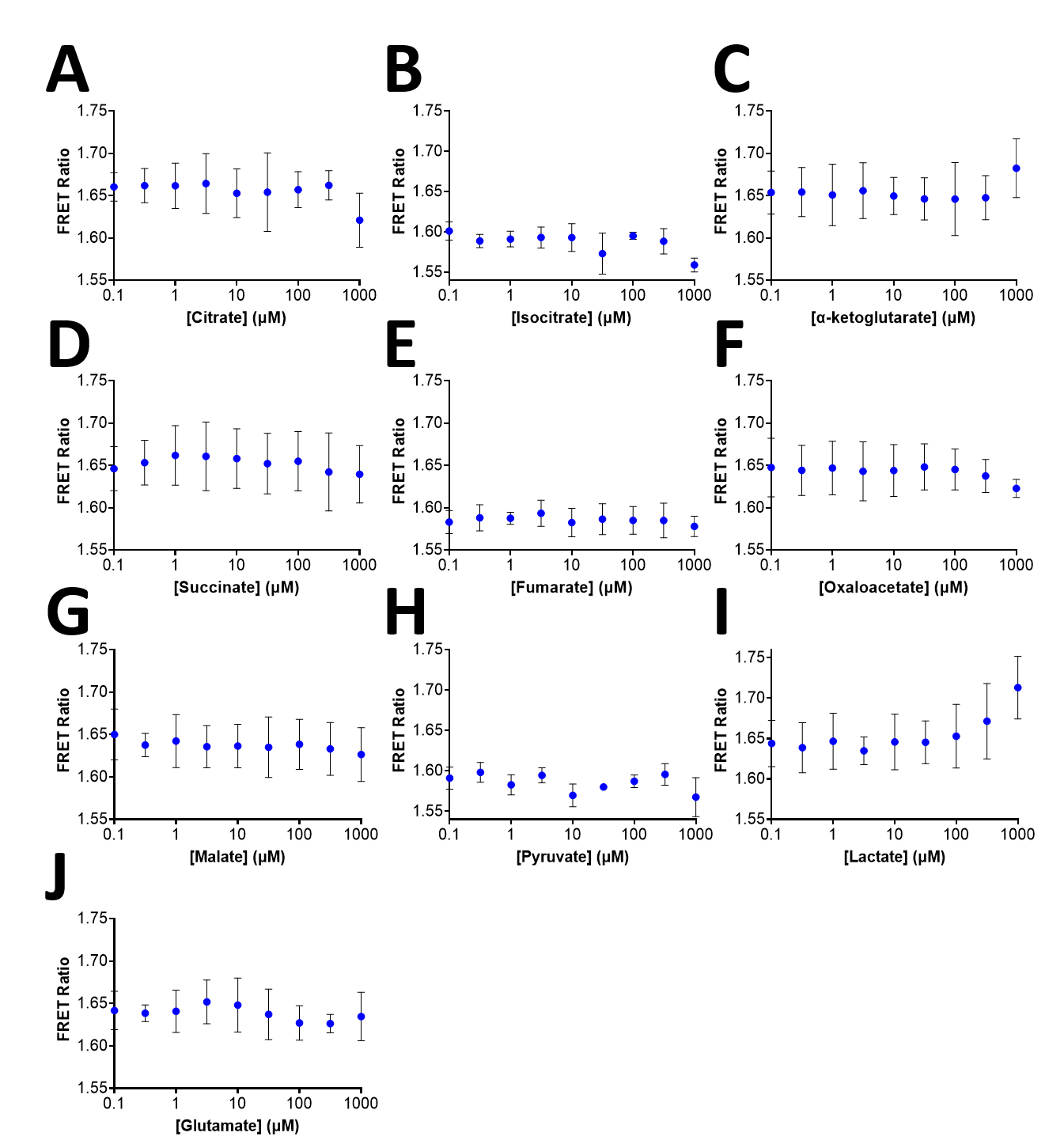

**Supplemental Figure 3. D2HGlo does not respond to Citric Acid Cycle intermediates, pyruvate, lactate and glutamate.** Purified sensor was titrated with increasing concentrations of: (A) Citrate, (B) Isocitrate, (C) α-ketoglutarate, (D) Succinate, (E) Oxaloacetate, (F) Malate, (G) Pyruvate, (H) Lactate and (I) Glutamate. The FRET ratio is plotted against the log concentration of each metabolite. Each data point represents the average ± standard deviation from three independent experiments.

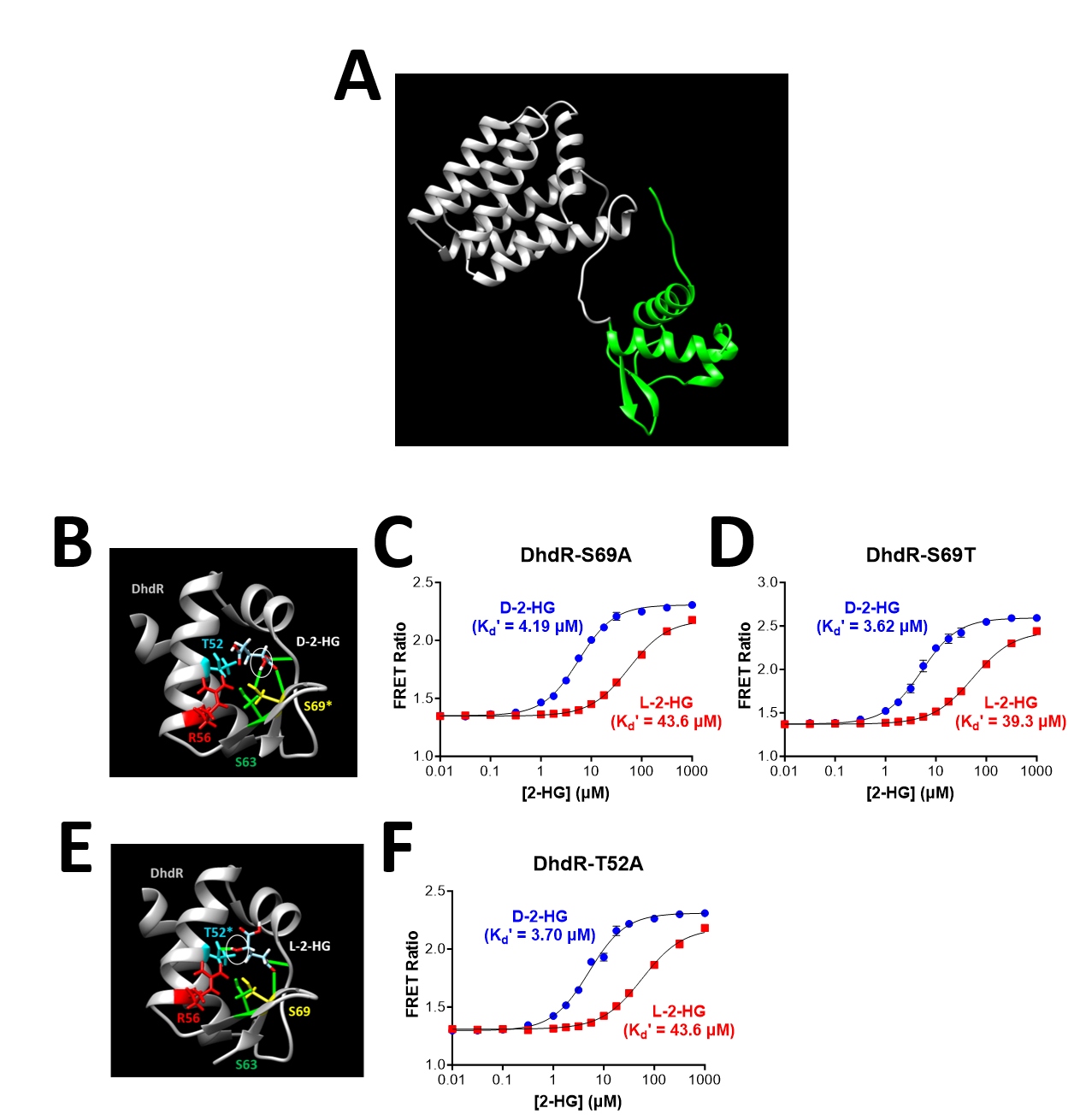

**Supplemental Figure 4. Site-directed mutagenesis of potential key residues in DhdR involved in D-2-HG or L-2-HG binding.** (A) 3D structure of DhdR generated in AlphaFold with the putative ligand binding domain shown in green. (B) Potential DhdR binding pocket for D-2-HG with putative residues involved in ligand binding (e.g. serine-69) highlighted. (C) 2-HG binding curve for the DhdR-S69A mutant. (D) 2-HG binding curve for the DhdR-S69T mutant. (E) Potential DhdR binding pocket for L-2-HG with putative residues involved in ligand binding (e.g. threonine-52) highlighted.. (F) 2-HG binding curve for the DhdR-T52A mutant.

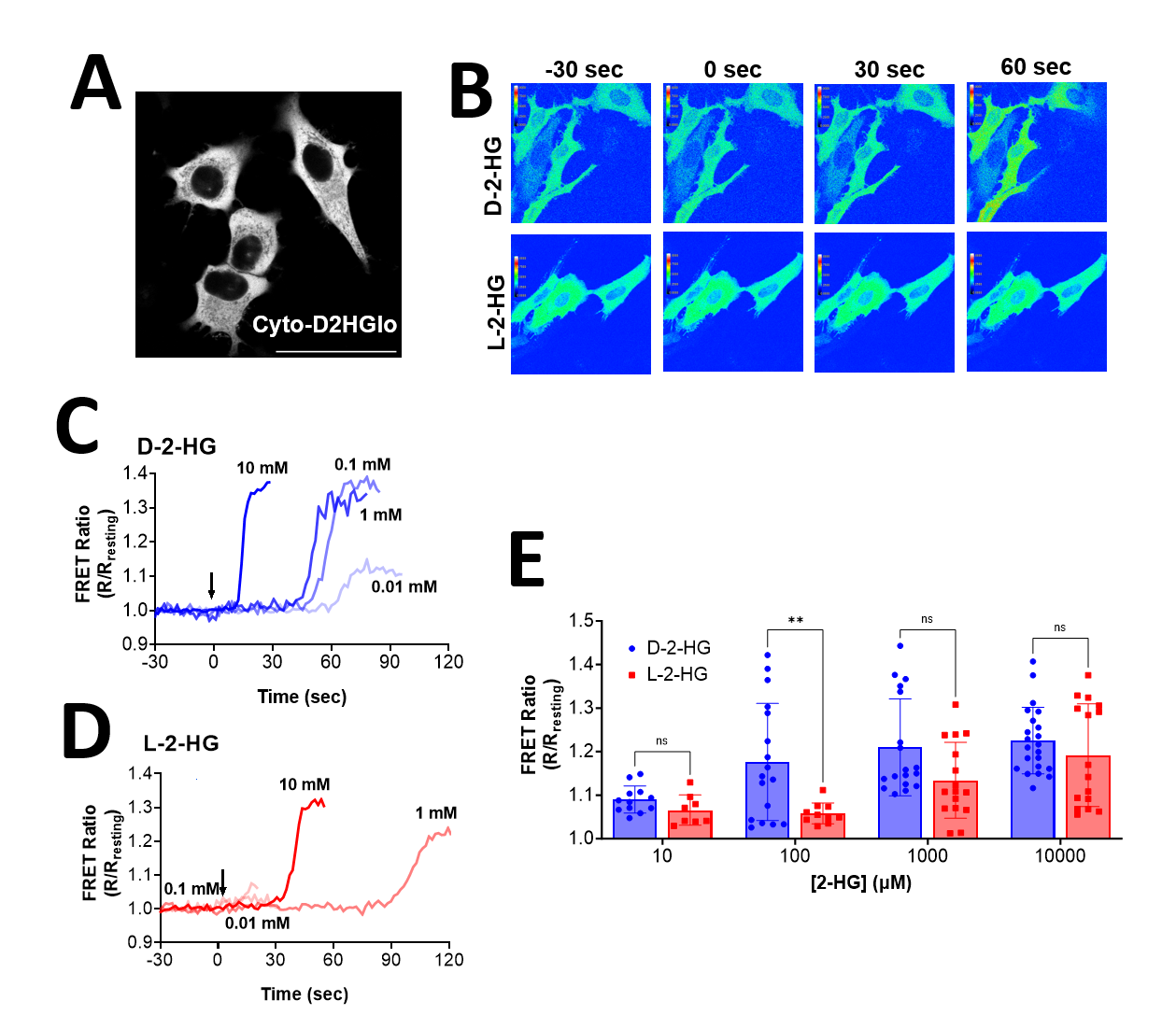

**Supplemental Figure 5.** **Imaging intracellular D-2-HG in HeLa cells using Cyto-D2HGlo.** (A) Representative image of HeLa cells expressing Cyto-D2HGlo. Scale bar is 50 µm. (B) Ratiometric FRET image of HeLa cells expressing Cyto-D2HGlo before and after addition of 1 mM D-2-HG (top) or 1 mM L-2-HG (bottom). Images are shown at 30 seconds before 2-HG addition (-30 seconds), at the time of addition (0 seconds) or for 30-60 seconds following addition. (C) Time-lapse imaging of HeLa cells expressing Cyto-D2HGlo following exposure to increasing concentrations of D-2-HG. The FRET ratio was collected for 30 seconds prior to the addition of digitonin (25 µM) and 0.01 mM, 0.1 mM, 1 mM or 10 mM D-2-HG. The FRET ratio (R) at each time point was normalized to the average FRET ratio recorded during the baseline measurements (R_resting_). (D) Time-lapse imaging of HeLa cells expressing Cyto-D2HGlo following exposure to increasing concentrations of L-2-HG. Data is presented the same as in panel C. (E) HeLa cells expressing Cyto-D2HGlo showed a concentration-dependent increase in the maximum FRET ratio that could be achieved with either D-2-HG and L-2-HG. Cells were treated with 25 µM digitonin and increasing concentrations of D-2-HG or L-2-HG, ranging from 0.01-10 mM. For D-2-HG, the bar graph represents n=12 cells from four independent experiments (0.01 mM D-2-HG), n=17 cells from six independent experiments (0.1 mM D-2-HG), n = 18 cells from six independent experiments (1 mM D-2-HG) and n=22 cells from eight independent experiments (10 mM D-2-HG). For L-2-HG, the bar graph represents n=8 cells from three independent experiments (0.01 mM L-2-HG), n=10 cells from five independent experiments (0.1 mM L-2-HG), n = 16 cells from six independent experiments (1 mM L-2-HG) and n=16 cells from five independent experiments (10 mM L-2-HG). Statistical analysis was performed using a two-way ANOVA test with *post hoc* Tukey (**, P < 0.01).

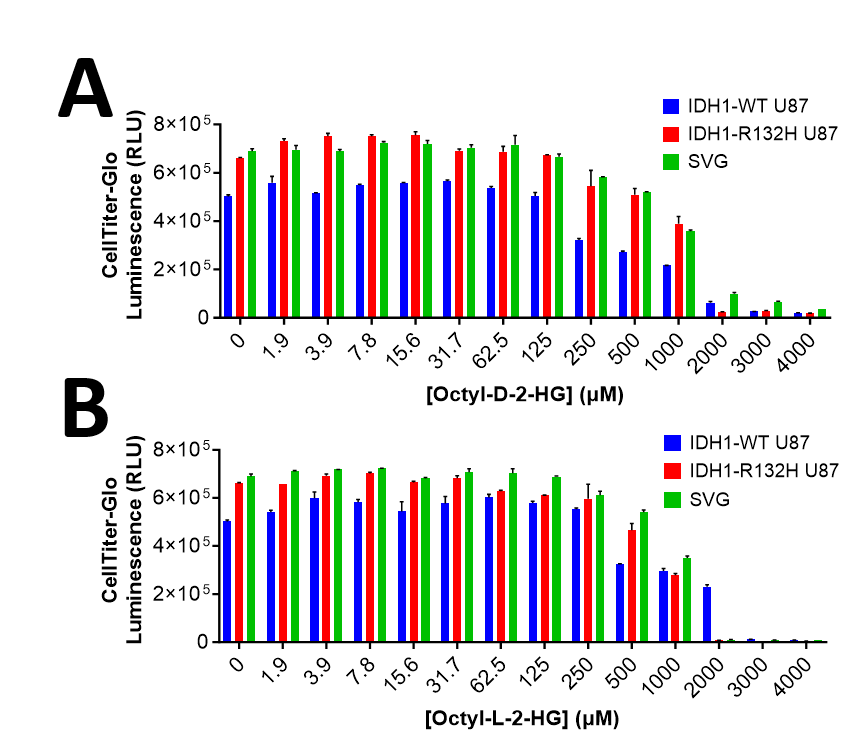

**Supplemental Figure 6. Viability of glial cells decrease following exposure to elevated concentrations of membrane-permeable D-2-HG or L-2-HG.** (A) Cell viability of *IDH1*-WT U87MG cells, *IDH1*-R132H mutant U87MG cells and human fetal glial SVG cells was measured using CellTiter-Glo® Luminescent Cell Viability Assay. Cells were allowed to attach overnight then treated for 48h with concentrations of octyl-D-2-HG ranging from 1.9 µM–4 mM. (B) Same set of experiments performed with octyl-L-2-HG.

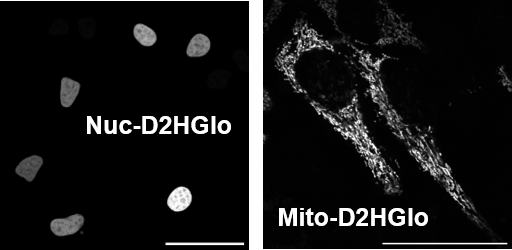

**Supplemental Figure 7. Expression of Nuc-D2HGlo and Mito-D2HGlo in HeLa cells.** HeLa

cells were transfected with Nuc-D2HGlo or Mito-D2HGlo and imaged on a confocal microscope

at 48 hours post-transfection. Scale bar is 50 μm.

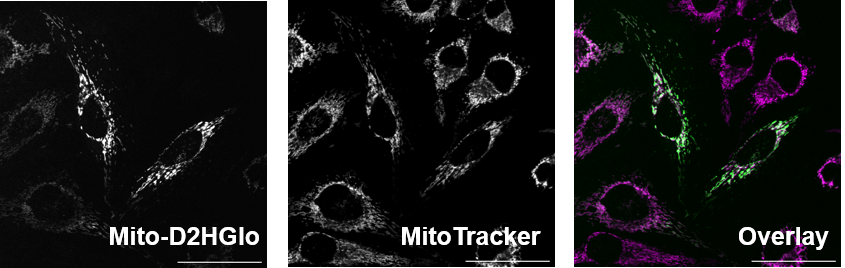

**Supplemental Figure 8. Colocalization of Mito-D2HGlo with MitoTracker DeepRed**. HeLa

cells were transfected with Mito-D2HGlo. At 48 hours post-transfection, cells were exposed to

MitoTracker for 10 min prior to fluorescence imaging. Separate images of Mito-D2HGlo and

MitoTracker DeepRed are shown. An overlay of both channels is shown on the right.

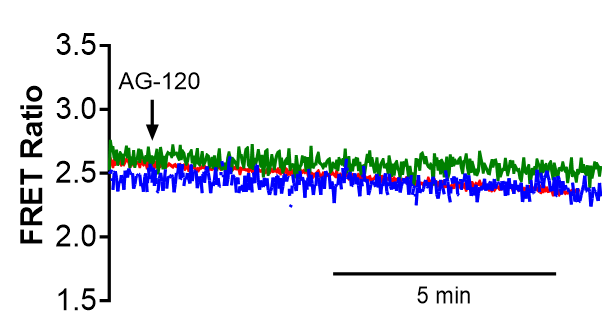

**Supplemental Figure 9. *IDH1*-R132H mutant U87MG cells expressing Cyto-D2HGlo do not respond to acute treatment with AG-120.** Cells expressing Cyto-D2HGlo (48h post-transfection) were treated with AG-120 (10 μM) and the FRET ratio was monitored for approximately 15 minutes after drug addition. Each line represents a single cell treated with AG-120.

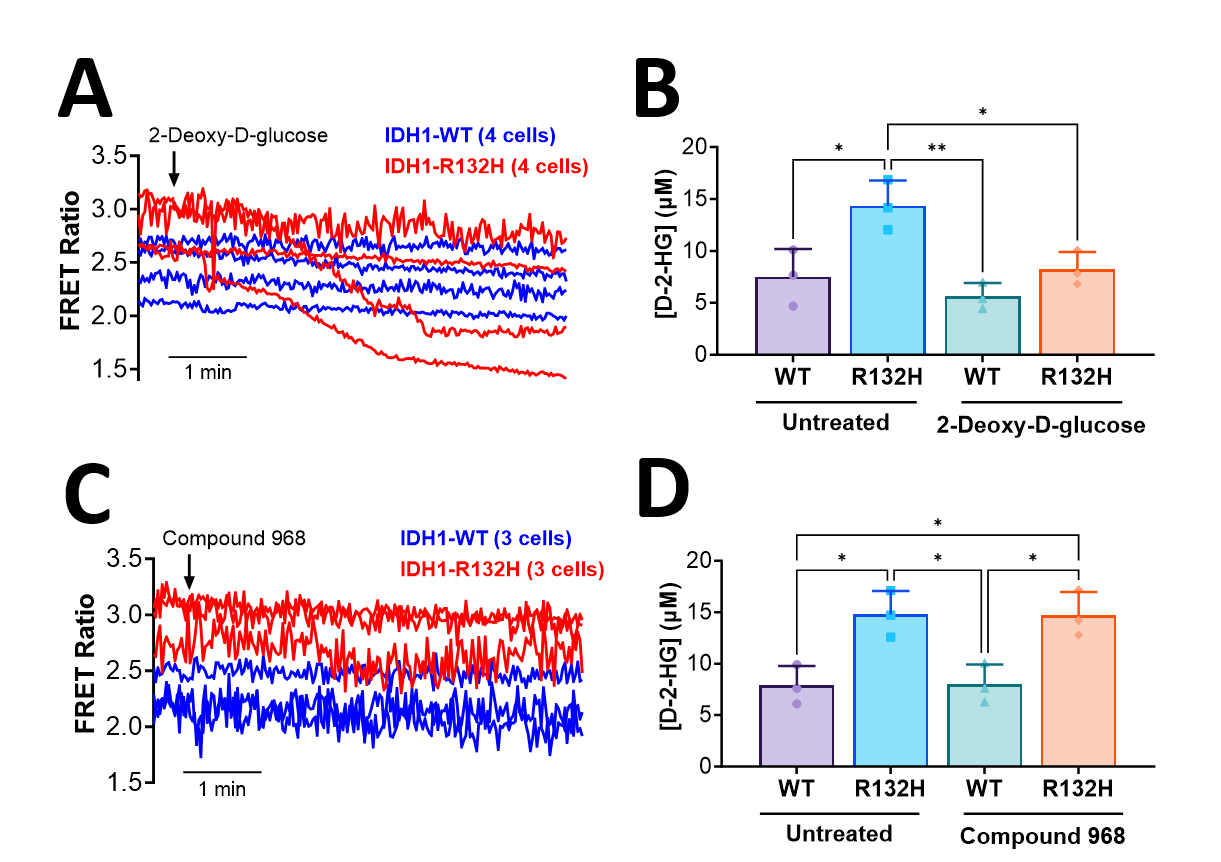

**Supplemental Figure 10. D-2-HG production is likely driven by glycolytic metabolism but not glutaminase in *IDH1*-R132H U87MG cells**. (A) *IDH1*-wildtype or *IDH1*-R132H mutant U87MG cells expessing Cyto-D2HGlo were treated with 10 mM 2-Deoxy-D-glucose (2-DG) and the FRET raito was monitored for approximately 5.5 minutes. Each line represents a single cell treated with 2-DG (n=4 cells for WT and R132H). (B) *IDH1*-wildtype or *IDH1*-R132H mutant U87MG cells were treated with 2-DG (10 mM) for 48 hours after which the concentration of D-2-HG was determined in culture supernatants. Statistical analysis was performed using a one-way ANOVA test with *post hoc* Tukey (**, P < 0.01; *, P < 0.05). (C) *IDH1*-wildtype or *IDH1*-R132H mutant U87MG cells expessing Cyto-D2HGlo were treated with 1 µM Compound 968 (glutaminase C inhibitor) and the FRET raito was monitored for approximately 5.5 minutes. Each line represents a single cell treated with Compound 968 (n=3 cells for WT and R132H). (B) *IDH1*-wildtype or *IDH1*-R132H mutant U87MG cells were treated with Compound 968 (1 µM) for 48 hours after which the level of D-2-HG was assessed in culture supernatants. Statistical analysis was performed using a one-way ANOVA test with *post hoc* Tukey (*, P < 0.05).

**Supplemental Table 1. *In vitro* characterization of D2HGlo at different pH and temperatures.** Values shown are the average of three independent experiments. The K_d_’ of D2HGlo remained unchanged across three different pH values. In contrast, the dynamic range was dramatically decreased at pH 6.5 and was increased at pH 8, relative to measurements performed at pH 7.4. The K_d_’ and dynamic range of D2HGlo are not drastically altered at 30ºC or 37ºC.

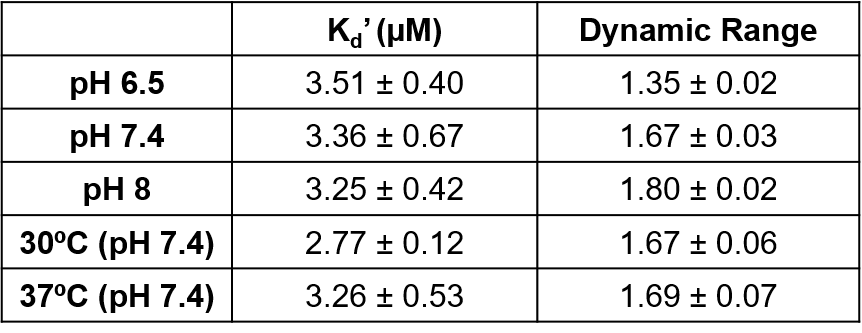

**Supplemental Table 2. Primers used in molecular cloning**. The forward and reverse primers for generating the initial DhdR-based sensor construct, three D2HGlo linker variants, three DhdR binding pocket mutants and the three cellular D2HGlo constructs.

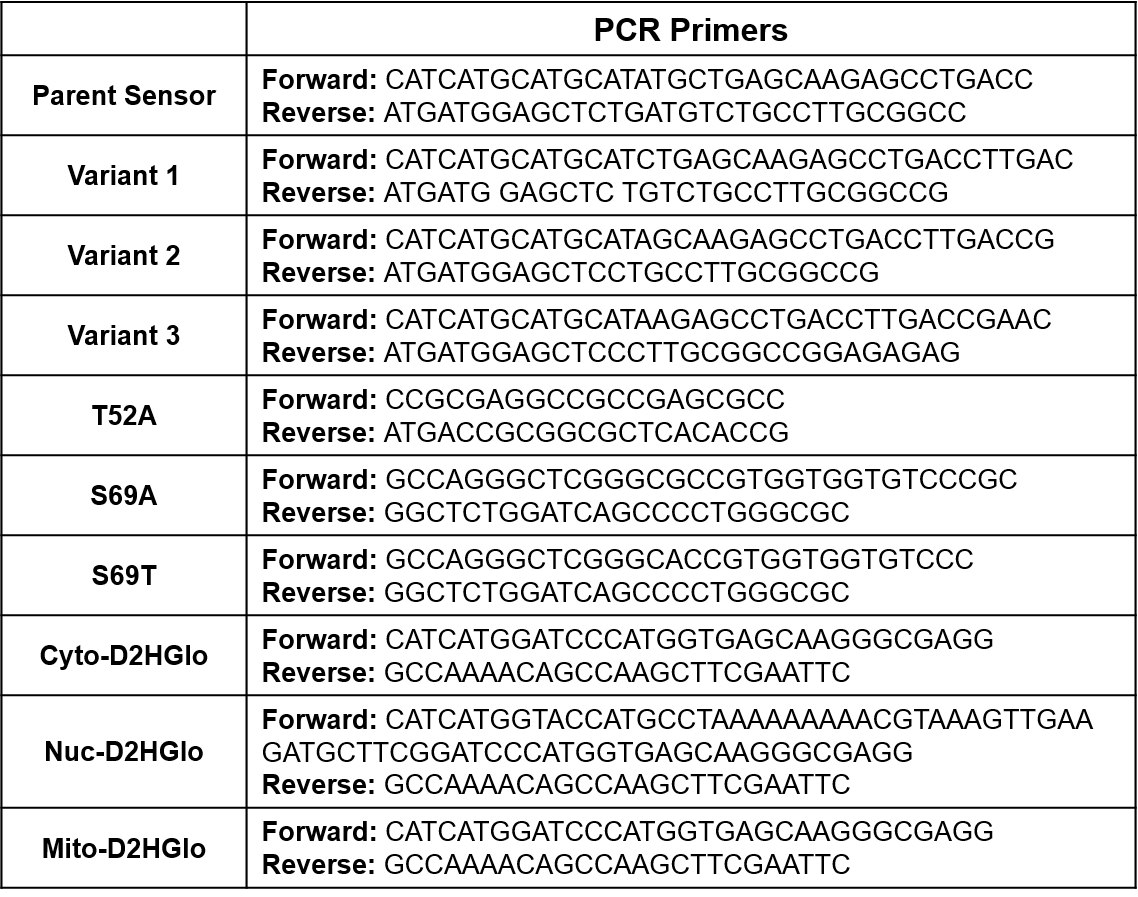
